# Ancient Somatosensory Circuit Architectures Employ Flexible Molecular Strategies

**DOI:** 10.64898/2026.08.25.747052

**Authors:** Hyemin Lee, Paul W. Frazel, Elena Singer-Freeman, Anne E. Cavanagh, Hanshin D. Shin, Kalina Rice, Shanmugapriya Selvaraj, Mark Alu, Heebal Kim, Cynthia Loomis, Shane A. Liddelow, Myungin Baek, Jeremy S. Dasen

## Abstract

The extent to which conserved neural circuit architectures depend on shared molecular specification programs remains unclear. Here, we address this question by examining the somatosensory system of the little skate, *Leucoraja erinacea*, an early-diverging vertebrate that retains ancestral features of both finned and limb-based body plans. We show that core features of somatosensory circuit organization, including laminar organization of the spinal cord and dorsally restricted targeting of sensory afferents, are deeply conserved. Unexpectedly, the molecular programs specifying dorsal root ganglion (DRG) sensory subtypes diverge extensively from those of mammals. Although DRG neuron subtype specification and spinal connectivity rely on target-derived cues, skates employ distinct neurotrophin receptor and transcription factor identity codes. These findings support a model in which conserved spinal circuit architectures provide a stable scaffold that leverages flexible sensory neuron specification programs, enabling the evolutionary diversification of vertebrate somatosensory systems.

## Introduction

Across bilaterians, nervous systems preserve core features of an ancestral patterning plan despite extensive diversification in the mechanisms governing fate specification and circuit assembly^1-3^. A central question is whether conserved neural circuit architectures depend on shared developmental programs. Addressing this problem requires comparative analyses in phylogenetically distant vertebrates capable of distinguishing ancestral features from derived innovations.

The vertebrate somatosensory system provides an informative model for examining the evolution of neural circuits, owing to the extensive diversification of sensory modalities across species. In mammals, somatosensation is mediated by over a dozen genetically and physiologically distinct dorsal root ganglion (DRG) neuron subtypes^4-9^. Three principal classes - mechanoreceptors, nociceptors, and proprioceptors – transmit modality-specific information to spatially organized circuits within the spinal cord^10-13^. During development, post-migratory neural crest cells coalesce adjacent to the spinal cord to form segmentally arrayed DRG^14^. In contrast to the central nervous system, where neural diversity is generated primarily through spatial and temporal patterning mechanisms^3,15-18^, early-born DRG neurons are initially transcriptionally similar^5^. As sensory axons extend into the periphery, target-derived signals initiate programs that give rise to modality-specific molecular features. Each of the three major classes of mammalian somatosensory neurons preferentially expresses one of three neurotrophin receptors – *ntrk1* (nociceptors), *ntrk2* (mechanoreceptors), and *ntrk3* (proprioceptors) (Figure 1A). Signaling through neurotrophin receptors regulates multiple aspects of DRG maturation, including molecular identity, central connectivity, and neuronal survival^17,19-22^. The evolutionary origin of this target-dependent strategy for circuit assembly remains unknown.

**Figure 1.**
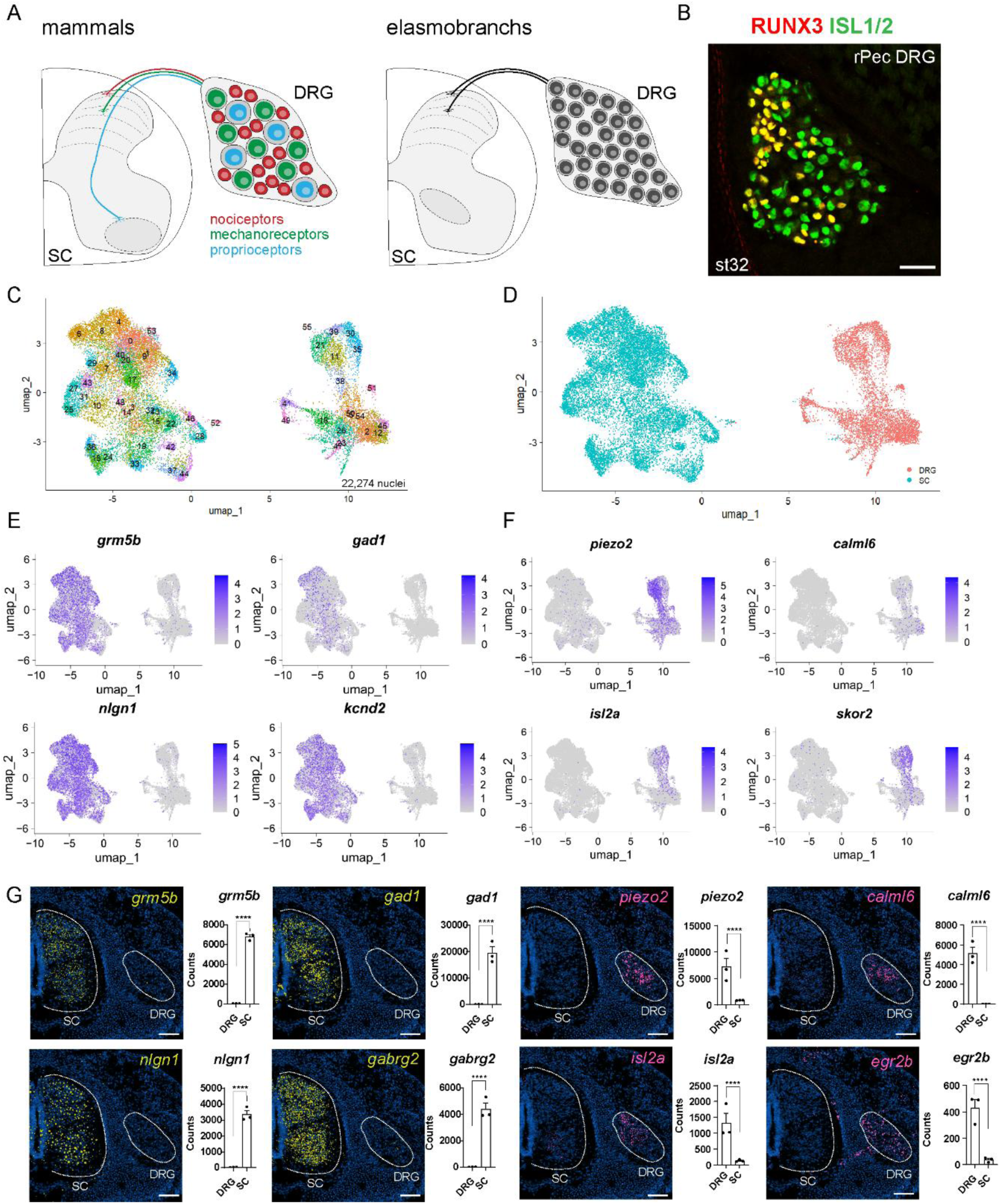
Integrated analyses of spinal cord (SC) and dorsal root ganglia (DRG) cells in *Leucoraja*. (A) Schematic showing organization of DRG classes in mammals and unknown types of elasmobranchs. Light grey shading of SNs indicated myelination. Gray oval in SC shows motor neuron position. (B) Expression of RUNX3 and ISL1 in skate rostral pectoral (rPec) DRG, indicating molecularly diverse subtypes. Scale bar, 50 µm. (C) Uniform manifold approximation and projection (UMAP) plots of nuclei from SC and DRG samples. Note DRG samples contain non-neuronal cells (D) Low resolution clustering showing transcriptional differences between SC and DRG samples. (E) Feature plots of select genes enriched in SC. (F) Feature plots of DRG-enriched genes. (G) Xenium spatial transcriptomic analyses of indicated genes enriched in SC and DRG. Images are shown from a single rostral pectoral section at st31. Scale bars, 100 µm. Similar patterns were observed in multiple segments from n=4 embryos analyzed at st31+ (n=2) and st32 (n=2). Plots on right show mean ± SEM RPKM from three biological replicates. Statistical significance was determined from raw read counts using edgeR with TMM normalization and a negative-binomial model, with Benjamini–Hochberg correction for multiple testing. ****FDR < 0.0001. See also Figures S1, S2, S3, S4 and Tables S1 and S2.

Early diverging vertebrate lineages provide an entry point for reconstructing the ancestral organization of somatosensory circuits. These species diverged prior to the radiation of tetrapods yet exhibit complex sensorimotor behaviors^23,24^. Prior studies of sensory systems in aquatic vertebrates have largely focused on specialized modalities including the lateral line, electrosensory organs, or Rohon-Beard neurons^25-29^, leaving the evolutionary history of canonical somatic sensations unexplored. Elasmobranchs (which include skates, sharks, and rays) are of particular interest as they represent an early vertebrate lineage in which primary somatosensory neurons are encapsulated in DRG^30^.

Anatomical and functional studies suggest that elasmobranchs possess a comparatively simplified somatosensory system, characterized by uniformly sized DRG soma and bimodal distributions of axonal diameters and stimulus response profiles^31,32^. Notably, elasmobranchs lack unmyelinated C-fibers, which transmit nociceptive signals and constitute the majority of DRG neurons in mammals (Figure 1A) ^31-36^. While neurons in the elasmobranch spinal cord are arranged in histologically distinct layers^37,38^, the composition of its somatosensory neuronal subtypes remains undefined. It is unknown whether elasmobranch DRG contain molecularly distinct nociceptor and mechanosensory populations, how such neurons are specified during development, or whether they engage conserved spinal circuits associated with pain and touch processing. This gap limits our ability to reconcile functional and behavioral studies of somatosensation in basal vertebrates with modern genetic models of somatosensory neuron _diversity11,35,36,39._

The little skate *Leucoraja erinacea* provides a tractable system for investigating the evolution of somatosensory diversity and circuit organization^23^. Skates employ a bipedal, walking-like fin locomotor behavior controlled by spinal neurons that share genetic signatures with those of tetrapods^40-42^. Whether this deep conservation of motor circuits extends to the somatosensory system remains unknown. Here, we integrate molecular profiling, spatial transcriptomics, and embryological perturbations to test whether conserved circuit architectures depend on shared molecular specification programs and to explore conserved mechanisms that generate sensory neuron diversity.

## Results

### Integrated molecular and spatial analysis of somatosensory neuronal diversity in *Leucoraja*

To assess molecular diversification of DRG neurons in *Leucoraja*, we examined the temporal emergence of sensory neuron (SN) fate determinants relative to fin development, a major peripheral target. We analyzed the expression of ISL1, a transcription factor broadly expressed in tetrapod DRG neurons^43-45^, and RUNX3, a factor essential for the development of mammalian proprioceptive SNs^46,47^. ISL1^+^ was detected in DRG neurons between the stages when fins form (st28) and become mobile (st32), whereas RUNX3 expression first appeared at st29 (Figure S1A-C). RUNX3 was detected in ∼45-70% of DRG neurons through st32, with a higher proportion of sensory neurons expressing RUNX3 at pelvic and tail segments (Figure 1B, S1D-E). In rostral pectoral segments, RUNX3 localized to the medial DRG, whereas in more caudal segments RUNX3^+^ neurons were intermingled with ISL1^+^, RUNX3^-^ SNs, resembling the intermixing of DRG subtypes observed in mammals (Figure 1B, S1F). These observations indicate that DRG neurons in *Leucoraja* exhibit early molecular heterogeneity despite limited anatomical segregation of sensory subtypes.

To further examine the molecular composition of DRG and spinal cord (SC) populations, we performed bulk and single-nucleus RNA sequencing (snRNAseq) on tissue collected from st31-st32 skate embryos. We chose st31-st32 embryos for our profiling in order to capture expression of developmental determinants that may be downregulated in mature spinal and DRG neurons. For bulk RNAseq, we prepared three biological replicates, each consisting of pooled spinal cord (SC) and DRG tissues from multiple (3-4) embryos and performed a differential expression analysis between the two tissue types (Figure S2A). For snRNAseq, we employed PIPseq, which enables direct capture of nuclei from frozen SC and DRG tissues^48,49^. We collected nuclei from pooled whole SC (6 embryos) and individually isolated DRGs (14 embryos) for each of two independent sequencing runs. After filtering out low-quality nuclei, datasets from both runs were merged, yielding a total of 22,724 nuclei (15,697 from SC and 6,577 from DRG) (Figure 1C, D).

Comparative analysis identified genes selectively enriched in SC or DRG populations (Figure 1E,F, S2A-C). Spinal cord nuclei were enriched for genes associated with inhibitory neurotransmission (e.g. *gad1*, *gabrg2*), consistent with an absence of inhibitory neurons in DRG (Figure 1E, G). DRG nuclei were enriched for genes associated with sensory modalities, including *piezo2* and *trp* channels^39,50^ (Figure 1F, S2C). We further identified transcription factors (TFs) selectively expressed by either SC or DRG tissues, including regulators of spinal progenitor fates (e.g. *olig2*, *nkx6.1*, *irx3*), postmitotic spinal neurons (*foxp2*, *lbx1, lhx5*), and DRG sensory neurons (*etv4*, *isl2*, *runx3*) defined in mammals (Figure S2B).

To validate these expression patterns and resolve their spatial organization, we performed 10x Genomics Xenium spatial transcriptomics on st31-st32 skate embryos across rostral pectoral (rPec), caudal pectoral (cPec), and pelvic (Pelv) segments (Figure S3A-B). We designed a custom 300-gene probe set from our profiling datasets enriched for cell fate determinants, signaling molecules, and known markers of SC and DRG subpopulations (Table S1). We prioritized cell fate determinants and genes likely to be critical for the emergence of subtype-specific functional properties, including channels and cell surface receptors. Approximately 90% of targeted genes yielded spatially informative expression patterns (Table S2). To further assess the fidelity of Xenium-based detection, we compared spatial expression patterns with quantitative expression data obtained from bulk RNA sequencing. Genes enriched by bulk RNA-seq showed largely concordant spatial localization in Xenium data, including SC-enriched genes (*grm5b*, *gad1*, *nlgn1*, *gabrg2*, *bcl11aa*, *pax2*) and DRG-enriched genes (*piezo2*, *calml6*, *tsapn8*, *frmpd1*, *isl2a*, *egr3b*) (Figure 1G, S4A-B). Together, these data provide a foundation for investigating somatosensory molecular diversity and organization in an early-diverging vertebrate.

### Deep conservation of laminar organization in the dorsal spinal cord of *Leucoraja*

Because mammalian sensory neurons target specific layers of the spinal cord, we next examined the diversity and potential laminar organization of the dorsal spinal cord in *Leucoraja*. Re-clustering of spinal nuclei identified 29 clusters, many of which exhibited molecular profiles consistent with mammalian subtypes (Figure 2A-C, S5, Table S4). For example, cluster 2 was enriched for known spinal motor neuron markers (*chat*, *mnx1*, *slit3*), whereas cluster 17 expressed markers characteristic of dorsal excitatory dI1 interneurons (*lhx2*, *lhx9*). To validate cluster assignments, we examined spatial expression of cluster-enriched genes using Xenium, enabling assignment of major classes including ventral and dorsal interneurons, spinal progenitors, floor plate cells, and glial populations (Figure 2C, S5). Graph-based clustering further revealed sharp architectural boundaries among broad spinal classes (Figure 2D, E).

**Figure 2.**
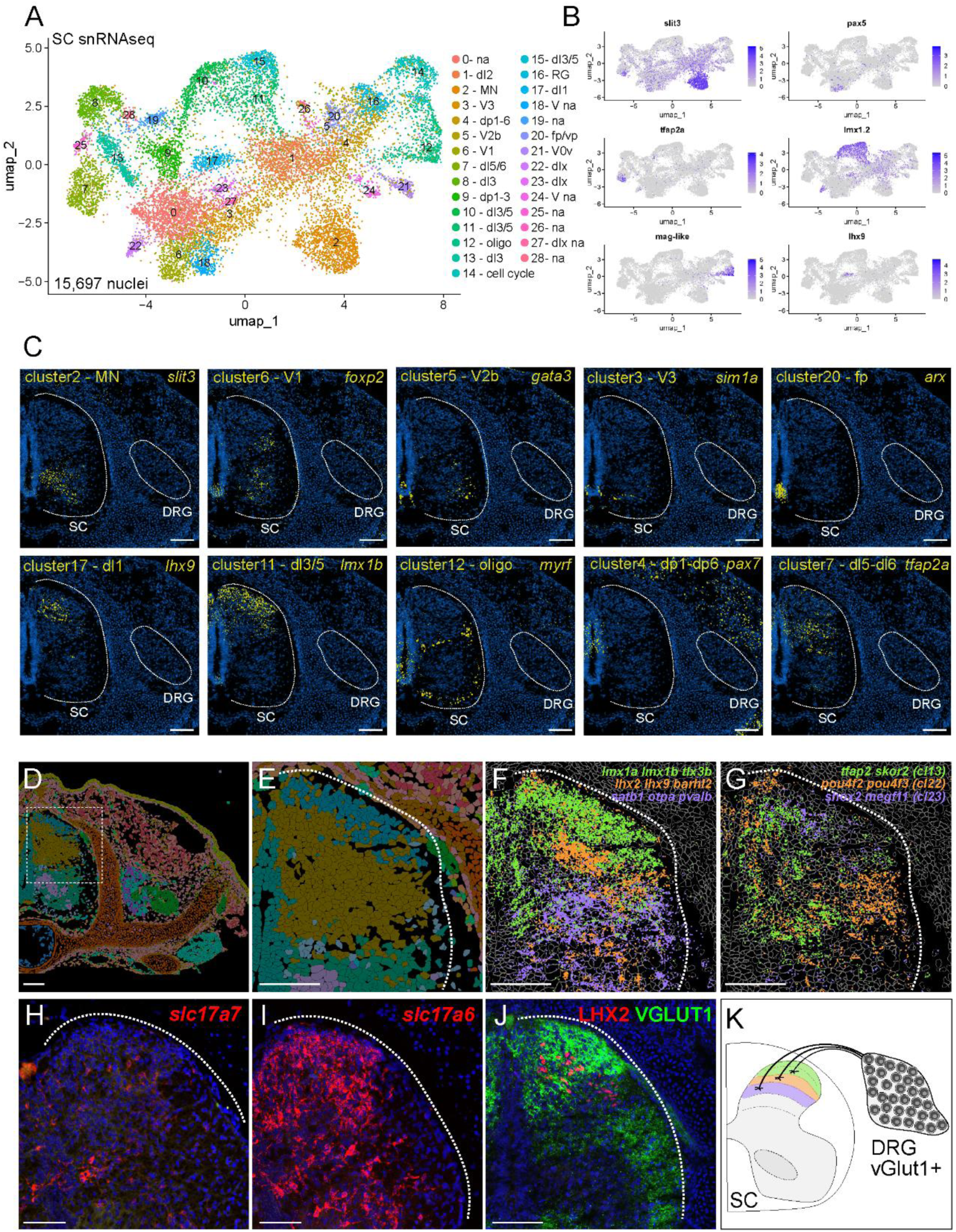
Diversity and laminar organization of spinal neurons in *Leucoraja*. (A) UMAP plot of reclustered SC nuclei. Putative homology to mammalian SC types based on expression patterns and conserved fate determinants. dp, dorsal progenitors; dI dorsal interneuron; dIx expressed in dorsal SC, V ventral SC, RG radial glia; p, progenitor, na, not assigned (B) Example feature plots of indicated cluster-restricted genes. (C) Xenium spatial transcriptomics of indicated cluster-restricted genes and assignment to cardinal spinal class. Images show transcript expression in yellow and DAPI stain in blue. Images are from a single section of rostral pectoral level at st31+. (D) Image of Xenium graph-based clustering of segmented cells in skate rostral pectoral section. (E) Magnification of image in panel D showing superficial and deep dorsal horn populations. (F) Expression of indicated genes in dorsal spinal cord. Expression for multiple genes are shown in a single color to highlight layering. (G) Expression of clustering-defining genes showing distinct populations along medial-lateral axis. (H-I) HCR in situ hybridization showing expression of *slc17a7* (*vglut1*) and *slc17a6* (*vglut2*) in st32 pectoral level spinal cord. DAPI in blue. (J) Staining of LHX2 and VGLUT1 in dorsal spinal cord of rPec levels at st32. (K) Model of VGLUT1^+^ inputs to layers of the dorsal SC. Scale bars, 50 µm. See also Figures S5 and S6.

We further examined the organization of neurons in the dorsal spinal cord. In mammals, dorsal spinal laminae receive modality-specific inputs: nociceptive afferents project predominantly to superficial laminae (I–II), low-threshold mechanoreceptive afferents in intermediate laminae (III-V), whereas proprioceptive afferents project to deeper dorsal laminae as well as ventral spinal circuits (Figure 1A) ^51,52^. To determine if a comparable organization exists in *Leucoraja*, we analyzed the spatial expression of conserved fate determinants. Neurons in the superficial dorsal horn expressed *lmx1a*, *lmx1b*, and *tlx3b*, markers of mammalian dI5 neurons^53,54^, that receive nociceptive input in mammals (Figure 2F). More ventral spinal neurons expressed *lhx2, lhx9,* and *barhl2*, markers of dI1 neurons that receive mechanosensory input^55,56^ (Figure 2F). Deeper spinal populations expressed *satb2*, *otpa,* and *palvb* (Figure 2F). Combinatorial analysis of cluster-restricted markers - including *tfap2*/*skor2* (cluster 13), *pou4f2*/*pou4f3* (cluster 22), and *shox2*/*megf11* (cluster 23) - identified interneuron subpopulations occupying stereotyped medial-lateral positions of the dorsal horn (Figure 2G).

To determine whether these layered spinal populations receive input from DRG sensory afferents, we leveraged conserved differential expression of glutamatergic markers in DRG and spinal cord. As in mammals, skate DRG neurons expressed both *vglut1* (*slc17a7*) and *vglut2* (*slc17a6*), whereas spinal neurons were largely devoid of *vglut1* expression (Figure 2H, I, S6A-D) ^57,58^. VGLUT1 can therefore be used to identify the central afferent projections of DRG neurons. We generated an antibody against VGLUT1 and examined the distribution of VGLUT1⁺ projections within the spinal cord. VGLUT1^+^ afferents terminals were enriched in the dorsal spinal cord and, by st32, extended toward the more ventrally positioned LHX2^+^ dI1-like spinal population (Figure 2J, S6E). VGLUT1 labeling was not detected in deeper layers of the ventral spinal cord, including near FOXP1 fin motor neurons (Figure S6F), consistent with prior DiI tracing studies of DRG-spinal projections in stingray^59^. Together, these data indicate that laminar organization of the spinal cord and dorsally restricted sensory inputs are deeply conserved features of vertebrate somatosensory systems (Figure 2K).

### Divergent neurotrophin receptor codes define sensory neuron identity in *Leucoraja*

While the profiles and topography of cardinal spinal neuronal classes show hallmarks of conservation at the molecular level, the diversity of elasmobranch DRG neurons is unknown. In mammals, at least 15 distinct subtypes of DRG sensory neurons have been described^9,10^. By contrast, anatomic and functional studies of elasmobranchs suggest a comparatively homogeneous population. Elasmobranch DRG soma are typically homogeneous in size, largely lack unmyelinated C-fibers, and display bimodal stimulus response profiles^32^. We therefore examined the molecular composition of DRG neurons to determine whether this apparent simplicity masks underlying molecular diversity.

While spinal nuclei consisted predominantly of neurons and glia, DRG samples contained non-neuronal cell types, including muscle, endothelial, and cartilaginous cells (Figure S7A,B), which we confirmed by spatial transcriptomics (Figure S7C). We used canonical sensory neuron markers (e.g. *runx3*, *ntrk2a*, *ntrk3*) to guide the selection of 3 clusters for reclustering, yielding seven clusters of presumptive DRG neurons (Figure 3A; S7B, S8C, Table S4). Spatial transcriptomic analysis of cluster-enriched genes confirmed localization within DRG neurons (Figure 3B, S8A). Consistent with both snRNAseq and mammalian profiles, medially positioned *runx3*⁺ neurons co-expressed *ntrk3* in rPec segments, whereas more lateral populations expressed *ntrk2a* (Figure 3B, S9A).

**Figure 3.**
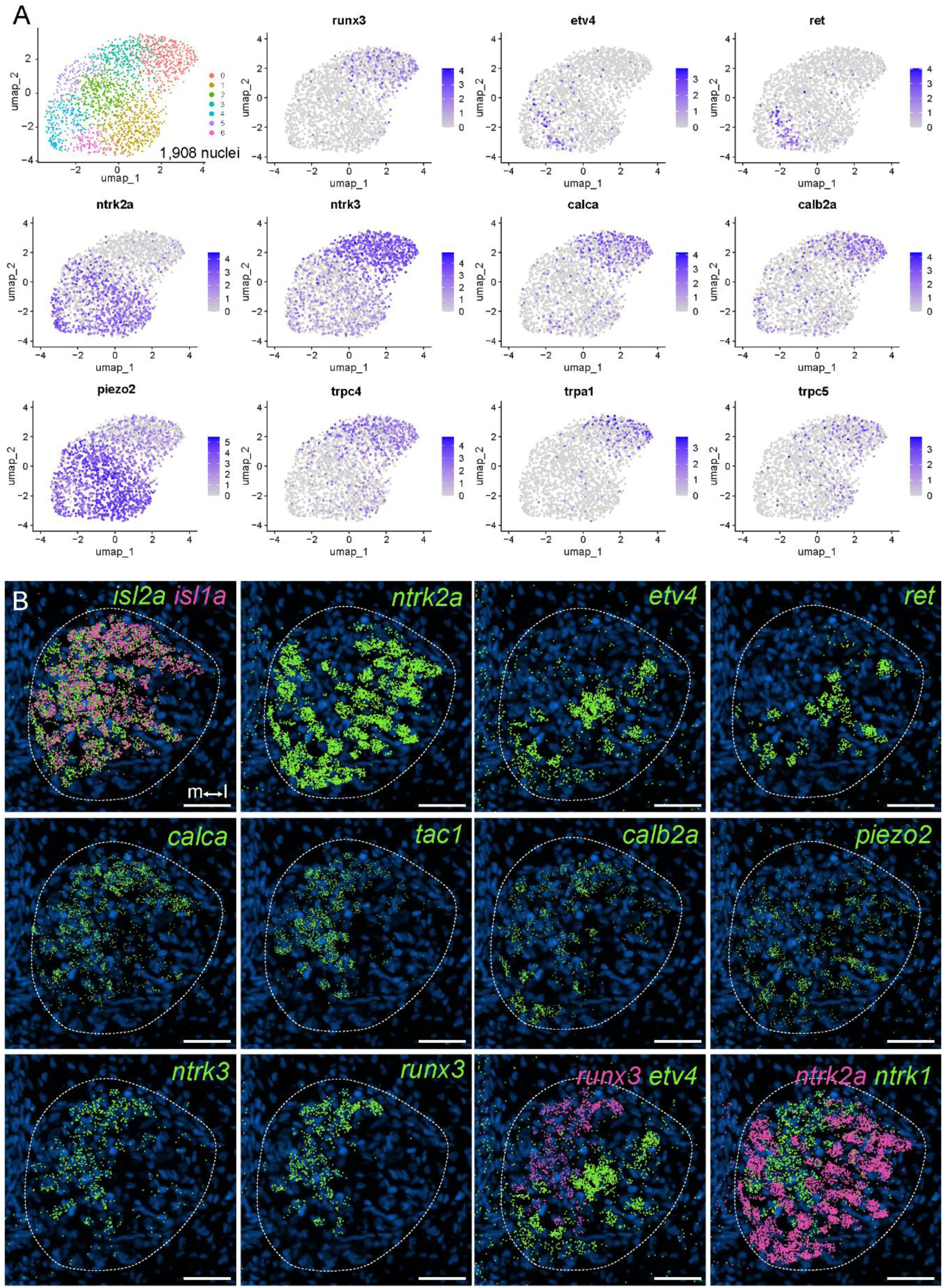
Molecular profile of DRG sensory neurons in *Leucoraja*. (A) UMAP and feature plots of reclustered DRG neurons. (B) Spatial transcriptomics of indicated genes in st31+ skate embryo at rPec segments. Expression of *isl1a* and *isl2a* shows DRG neuron position. Expression of *etv4* and *runx3*, as well as *ntrk1* and *ntrk2a*, are mutually exclusive. Medial (m) and lateral (l) positions are shown in panel B. Scale bars, 50 µm. See also Figures S7, S8 and Table S4.

Unexpectedly, *runx3^+^/ntrk3^+^* neurons, which mark muscle-innervating proprioceptive sensory neurons in mammals, co-expressed markers associated with nociceptive identity in mammals, including *trp* channels (e.g. *trpa1*) and *tac1* (Figure S9B-C). In addition, the observation that most skate DRG neurons could be classified by expression of either *ntrk2a* or *ntrk3*, was surprising given that mammalian nociceptors, which are typically *ntrk1^+^*, lack *ntrk3* expression. This prompted us to examine whether *ntrk1* was expressed in skate DRG. Although we were unable to detect *ntrk1* in our snRNAseq data (Figure S8B), likely due to the absence of *ntrk1* 3′-UTR sequence in the current skate genome annotation, spatial Xenium analyses revealed that *ntrk1* was expressed in *ntrk3*^+^ neurons (Figure S9A). We confirmed these observations using hybridization chain reaction (HCR) fluorescent *in situ* hybridization. These analyses revealed that *ntrk2a* was largely excluded from *ntrk3^+^* neurons, and most *ntrk3^+^* neurons co-expressed *ntrk1* (Figure 4A-B).

**Figure 4.**
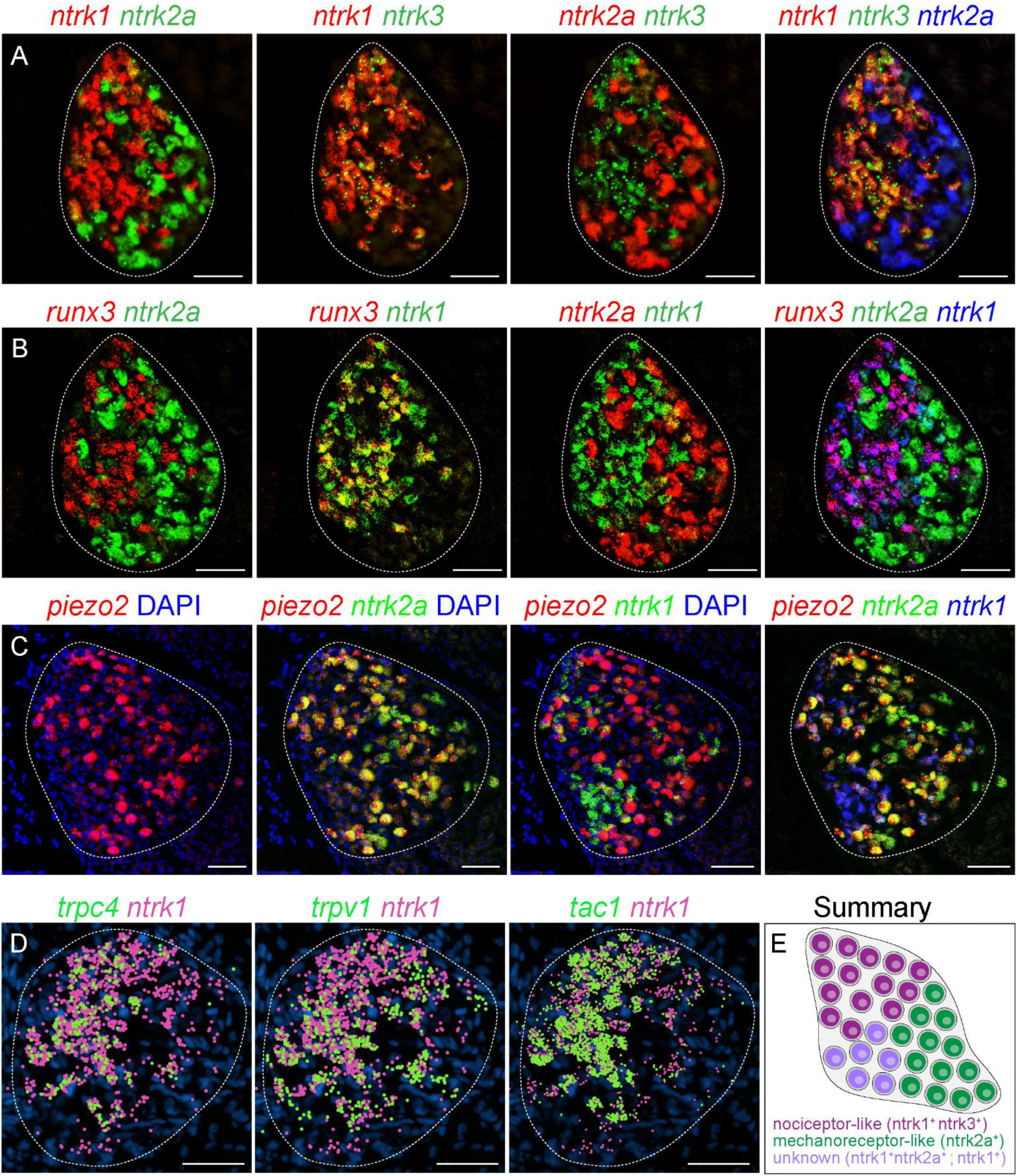
Divergent neurotrophin receptor codes in *Leucoraja.* (A) Expression of *ntrk1*, *ntrk2a*, and *ntrk3*. HCR *in situ* hybridization was performed on st31 skate DRG in rostral pectoral segments. (B) Expression of *runx3* in relation to *ntrk* gene expression at in rPec segments at st31. (C) HCR showing expression of *piezo2* in relation to *ntrk* gene expression at st32 in rPec segments. DAPI in blue. (D) Xenium spatial transcriptomic analyses of *trpc4*, *trpv1*, and *tac1* in relation to *ntrk1* expression in rPec segments at st31+. Scale bars, 50 µm. (E) Summary of putative DRG sensory classes in *Leucoraja*. See also Figure S9.

We quantified the distribution DRG neurons expressing combinations of *ntrk1*, *ntrk2a*, *ntrk3*, and *runx3* from our Xenium dataset. Although this approach has limitations, due to imprecise segmentation of cellular boundaries, they indicated patterns consistent with those observed by traditional histological methods and snRNAseq. Most *ntrk3^+^* neurons expressed *ntrk1* and *runx3*, and lacked *ntrk2a*, while *ntrk1^+^*cells consisted of those co-expressing either *ntrk2a* or *ntrk3*, or *ntrk1* alone (Figure S9F). A fourth population expressed *ntrk2*a and lacked *ntrk1* and *ntrk3* expression (Figure S9F).

Because *ntrk* expression often segregates with DRG neuronal classes in mammals, we further analyzed *ntrk1*, *ntrk2a*, and *ntrk3* in relation to functional determinants of sensory modality by both HCR and Xenium. DRG neurons expressing the mechanoreceptor channel *piezo2* expressed *ntrk2a*, consistent with expression of *ntrk2* as a conserved marker of mechanosensory DRG neurons (Figure 4C, S9E). We analyzed 4 of the *trp* channels we detected by bulk and snRNAseq (*trpv1*, *trpa1*, *trpm2*, and *trpc4*). *Trpv1* and *trpc4* were detected in *ntrk1/3*^+^ cells while *trpa1* was detected in a smaller subset (Figure 4D, S9B). Additional markers of mammalian C-fiber SNs, including *tac1*, were also detected in *ntrk1/3*^+^ cells (Figure 4D, S9C). Moreover, markers of recipient spinal neurons of nociceptors, such as *npy1r*, were enriched in the superficial layers of skate spinal cord (Figure S9D).

Together, these results demonstrate that sensory neuron identities in *Leucoraja* are defined by a divergent neurotrophin receptor code, in which molecular features associated with sensory modalities are combined within individual neurons (Figure 4E). These findings reveal that, despite a grossly conserved spinal circuit architecture, the molecular logic of sensory neuron specification has been extensively reconfigured during vertebrate evolution.

### Transcription factor expression delineates DRG sensory neuron subtypes

We next asked whether additional DRG subtypes are present within the broader sensory populations defined by *ntrk* expression. In many tissues, expression of cell fate determinants enables discrimination of distinct subtypes within a broader cellular class. Our molecular and spatial transcriptomic panel included 110 TFs, 32 of which were enriched in DRG cells (Table S3). Analyses of these TFs revealed four general categories of expression in DRG. The first class represents broadly expressed TFs across DRG neurons (e.g. *millt11b*, *tlx3b, pou4f3, six1*) (Figure 5A, S10A, F). The second and third classes of TFs were predominantly expressed in either *ntrk1/3^+^* neurons (*runx3*, *skor2*, *ebf3a, casz1*) or in *ntrk2^+^* neurons (*tbx2b, onecut2, etv4)* (Figure 5B,C, S10B,C,H). A fourth class included TFs that were expressed in overlapping subsets of *ntrk2* and *ntrk1/3* neurons (*shox2, phf24, runx1*) (Figure S10D,E,G). Together, these patterns suggest additional layers of neuronal diversity within the broader *ntrk1/3*⁺ and *ntrk2a*⁺ populations. For example, *etv4* and *onecut2* mark non-overlapping populations of *ntrk2a^+^* neurons (Figure 5C), suggesting a presence of distinct mechanosensory-like subtypes within *ntrk2a*⁺/*piezo2*⁺ populations.

**Figure 5.**
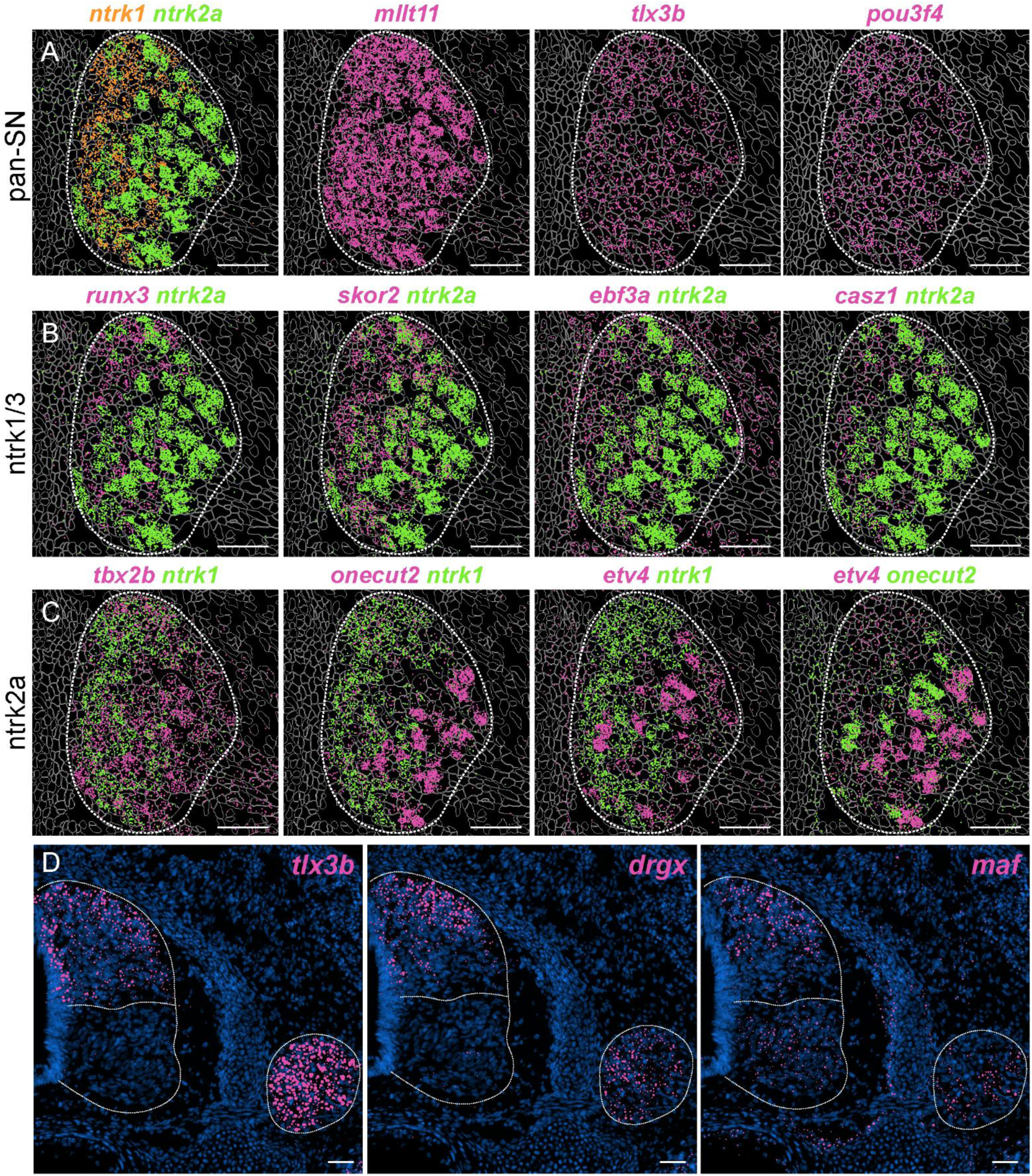
Delineation of DRG sensory subtypes by transcription factor expression. (A) Spatial transcriptomic analyses showing expression of *mllt11, tlx3b*, and *pou4f3*. (B) Expression of *runx3*, *skor2*, *ebf3a*, and *casz1* are enriched in *ntrk1/3*^+^ DRG cells. (C) Expression of *tbx2b*, *onecut2*, and *etv4* are enriched in *ntrk2a*^+^ cells. In panels A-C segmentation of cells is shown. (D) Expression of *tlx3b*, *drgx*, and *maf* are detected in both DRG cells and dorsal spinal cord. DAPI is shown in blue. Images from rPec segments at st31+. Scale bars, 50 µm. A subset of TF genes was also confirmed by HCR (Figure S10F-H). See also Figure S10 and Table S3.

Interestingly, several TFs expressed by DRG sensory neurons were also expressed by specific neuronal populations in the dorsal spinal cord. We identified nine TFs that were expressed by both DRG and dorsal SC neurons (*bcl11ba*, *drgx, maf, hey, pou4f2, pou4f3, shox2, skor2,* and *tlx3b*) (Table S3). Expression of *tlx3b* is broadly expressed in DRG and the dorsal spinal cord, while *drgx* and *maf* were detected in subsets of DRG and dorsal SC populations (Figure 5D). Coordinated expression of shared TFs in spinal and DRG subtypes may contribute to wiring specificity between connected neuronal classes, as proposed in other systems^60-62^.

### Ancestral role of target-derived cues in sensory and motor neuron development

The divergence of neurotrophin receptor and TF expression in skates raises the question of whether DRG neurons develop through mechanisms analogous to those of tetrapods. In mammals, target-derived cues play critical roles in shaping somatosensory circuit maturation by regulating neuronal identity, connectivity, and survival^17,19^. In the spinal cord, limb-derived signals direct the specification and organization of motor neuron (MN) pools^63-65^. In DRG neurons, target-derived cues promote the segregation of sensory classes from an initially transcriptionally unspecialized state and regulate central projection patterns (Figure 6A)^5,21,22^. By contrast, the few studies that have been performed in fish suggest a reduced or absent contribution of peripheral cues to motor circuit development^66^.

**Figure 6.**
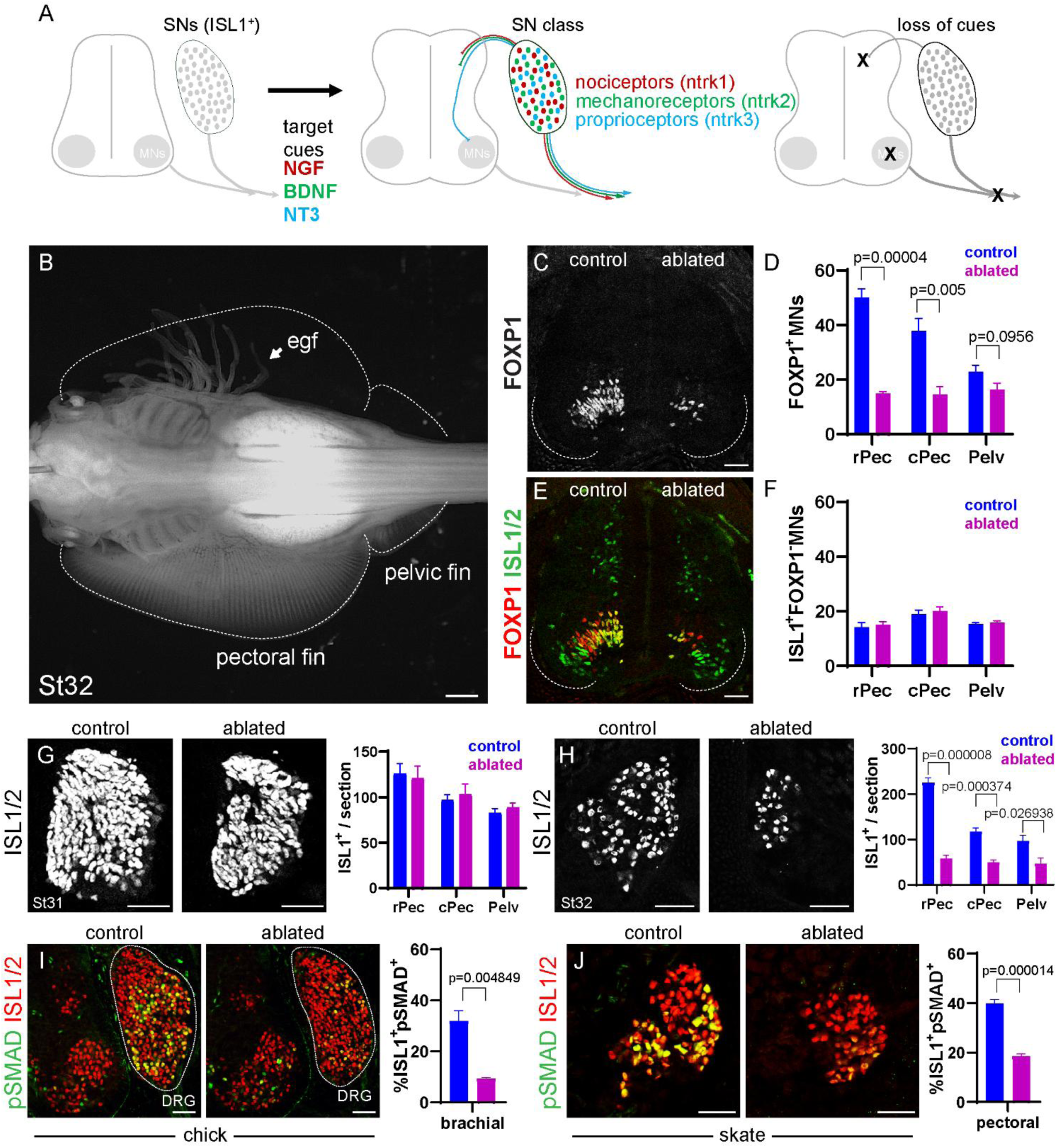
Role of fin-derived cues in somatosensory neuron development. (A) Model of maturation of DRG neurons through target-derived cues in mammals. Nerve growth factor (NGF), brain-derived neurotrophic factor (BDNF), and neurotrophin-3 (NT3) bind ntrk1, ntrk2, and ntrk3 respectively. In the absence of targets, SNs fail to mature and form appropriate central connections. (B) Example of fin-ablated embryo at st32 showing absence of fins on operated side. External gill filaments (egf) are exposed on ablated side. Scale bar, 1 mm. (C) Staining for the fin MN marker FOXP1. (D) Quantification of FOXP1 MNs in rostral pectoral (rPec), caudal pectoral (cPec), and pelvic (Pelv) segments. (E-F) Staining and quantification of FOXP1 and ISL1/2 neurons. (G) Staining and quantification of ISL1^+^ DRG neurons in control and ablated embryos at st31. (H) Staining and quantification of ISL1 in control and ablated embryos at st32. Neuron counts in H used Xenium samples. (I) Effect of chick forelimb bud ablation on SMAD phosphorylation. Graph on right shows percentage of SMAD^+^ ISL1/2^+^ neurons in brachial DRG. Images of ablated sides have been vertically inverted for clearer comparison. (J) Effect of fin bud ablation on SMAD phosphorylation in skate. Graph on right shows percentage of SMAD^+^ ISL1/2^+^ cells in pectoral DRG. For graphs in panels D, F, G-J cell counts per section were averaged from 4-8 sections/embryo in at least n=3 embryos. Bar graphs show the mean of the per-embryo averages ± S.E.M. Exact p-values shown when significant (unpaired t-test)., Scale bars in C-J, 50 µm. See also Figure S11.

To investigate the role of target-derived cues in sensorimotor circuit development in skates, we performed unilateral fin-ablation experiments and assessed the maturation of spinal and DRG neurons. Fin buds were ablated unilaterally near the time of their emergence (st27-st28) and allowed to develop until st31-st32 (14-28 days). In pilot experiments ∼90% of embryos survived and developed normally after combined unilateral ablation of the nascent pectoral and pelvic fin buds, except for the absence of fins on the operated side (Figure 6B).

We first examined the impact of fin removal on spinal MNs, which undergo rapid degeneration following limb bud ablation in chick embryos^67,68^. At st31 (12-17d post ablation), 65-70% of pectoral and 30-40% of pelvic MNs were lost, as assessed by reduced numbers of FOXP1^+^ neurons, a marker of tetrapod limb-innervating MNs^69-72^ (Figure 6C, D). By contrast, the number of ISL1^+^, FOXP1^-^ neurons, largely comprising axial-muscle innervating MN subtypes, was unaffected (Figure 6E, F). Consistent with selective vulnerability, spatial transcriptomic analyses revealed loss of several MN-restricted markers, including *chat*, *aldh1a2*, *chrnb4*, *mnx1*, *isl2a*, *tac1*, and *grfra4,* whereas markers of ventral (*foxp2*) and dorsal spinal interneurons (*lhx5*) were largely unaffected (Figure S11A).

To assess the impact of fin ablation on general features of DRG neurons, we analyzed expression of ISL1, as it is broadly expressed by embryonic DRG neurons and is essential for their specification^43^. In contrast to MNs, DRG neurons were relatively spared at early stages after fin ablation, consistent with studies in chick embryos^63,67^. At st31, fin ablation had little impact on DRG neuron number, as assessed by ISL1⁺ cells (Figure 6G). By st32, however, sensory neuron numbers were markedly reduced on the ablated side, with the most pronounced losses observed in rostral pectoral segments, where DRG neuron numbers declined by more than 70% (Figure 6H).

### Conserved SMAD activation marks early DRG responses to target-derived cues

The distinct temporal sensitivity of DRG neurons to peripheral deprivation likely reflects conserved differences in neurotrophic requirements between these populations^73^. To understand the role of target-derived cues in somatosensory circuit maturation, however, it is essential to determine when DRG neurons normally respond to peripheral signals. Because there are few markers that report loss of target-derived signals, the precise stage in which DRG neurons are affected is unclear. In rodent trigeminal ganglia, target-derived BMP signaling plays a critical role in SN specification, where it induces phosphorylation of SMAD1/5/8 (pSMAD) in ophthalmic subtypes^74^. In DRG neurons, BMP also induces pSMAD and cooperates with neurotrophin signaling to regulate peripheral innervation pattern^75^.

To assess whether SMAD phosphorylation is a conserved early marker of SNs responding to peripheral cues, we examined pSMAD in both chick and skate DRG. In chick, pSMAD was detected in brachial (forelimb-innervating) DRG by st27 where it labeled ∼30% of ISL1^+^ SNs (Figure 6I). In skate, pSMAD labeling was observed as early as st30, and was enriched in RUNX3^-^ neurons at fin levels, indicating preferential expression in *ntrk2a*^+^ mechanosensory-like subtypes (Figure S11B, C). In both species, pSMAD is first detected at stages corresponding to when sensory axons invade appendicular muscle^40^, consistent with regulation by peripheral cues.

We performed unilateral limb and fin bud ablation experiments to assess whether SMAD phosphorylation depends on target cues. Because skates develop ∼5 times slower than chick, we analyzed stages corresponding to the most mature stage after ablation prior to sensory neuron cell death (i.e. +3 days/st27 in chick and +14-18 days/st31 in skate)^67^. After limb ablation in chick, we found that the level of SMAD phosphorylation was reduced by ∼70% on the ablated side (Figure 6I). In skate embryos ablated at st28, pSMAD staining was diminished by ∼53% at st31 (Figure 6J). Collectively, these findings identify SMAD phosphorylation as a conserved early marker of activation of SNs by limb- and fin-derived signals.

### Target-derived cues couple sensory neuron identity with central connectivity in vertebrates

Having identified an early stage when DRG neurons respond to fin-derived cues, we next sought to determine how these signals contribute to SN specification. We first asked whether the initial segregation of *ntrk2a* and *ntrk1/3-*expressing neurons is altered by fin removal. At st31, *ntrk2a* and *ntrk1* expression remained segregated into medial-lateral populations in both control and fin-ablated embryos (Figure 7A, S12A,B), indicating that early neurotrophin receptor patterning is largely target independent.

**Figure 7.**
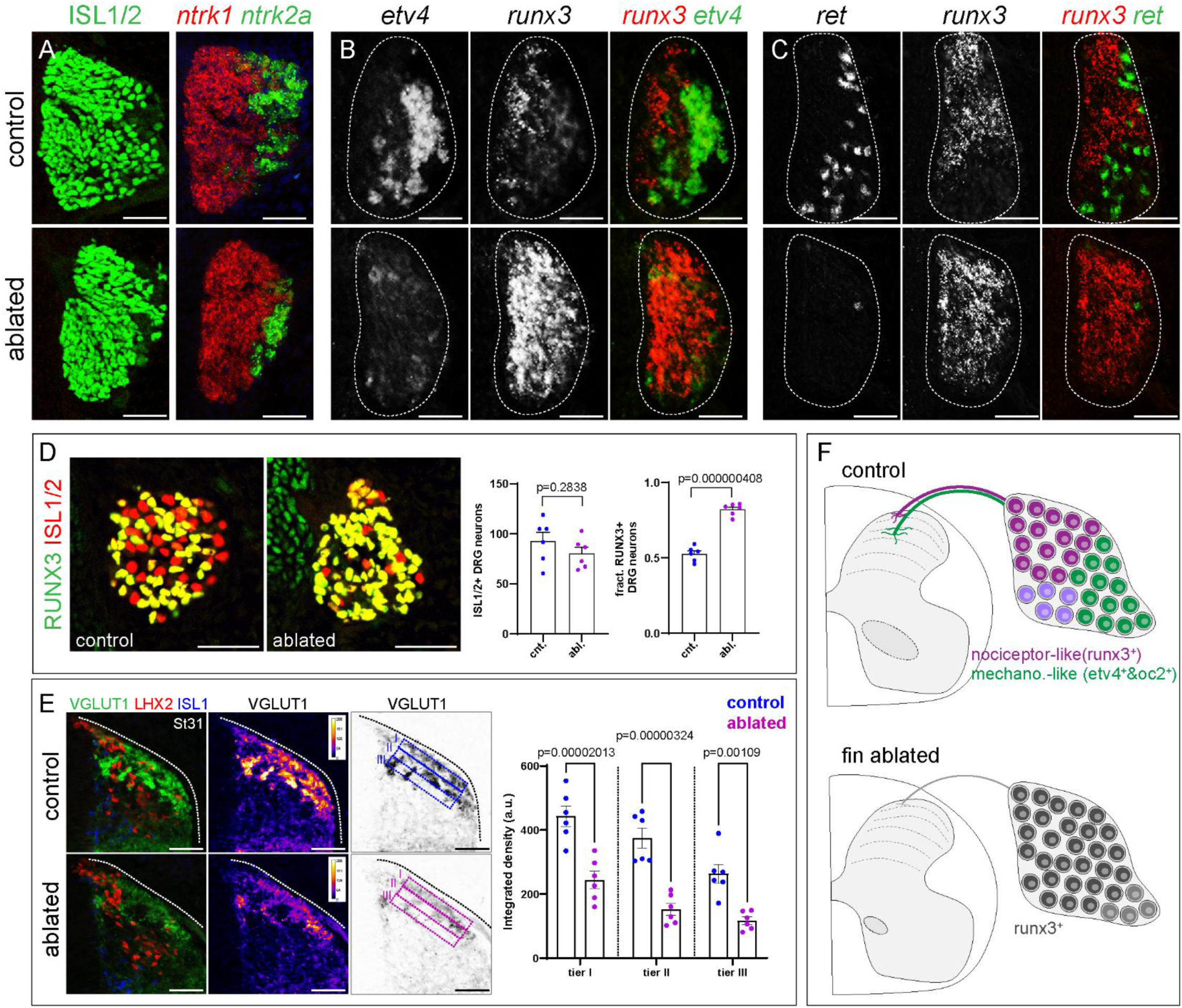
Target-derived cues shape DRG specification and central connectivity. (A) ISL1/2 staining (left) and *ntrk1*, *ntrk2a* HCR (right) on adjacent section of control and fin-ablated DRG. (B) Effect of fin ablation on *etv4* and *runx3* expression. (C) Effect of fin ablation on *ret* and *runx3* expression. (D) Quantification of RUNX3^+^ISL1/2^+^ neurons in control and fin-ablated embryos. Counts are averaged from n=6 embryos (4-7 rPec sections/embryo) at st31. Bar graphs show averages +/- S.E.M. Exact p values shown, unpaired t-test. (E) Reduction in VGLUT1 terminals after fin bud ablation. Plots on right show reduced signal density in three (I-III) regions of interest (ROI) averaged from n=6 embryos (5-6 rPec sections/embryo) at st31. Data are shown as mean ± SEM, with averages from individual embryos plotted on graphs. Control and Ablated groups were compared within each tier using two-way ANOVA followed by Šídák’s multiple-comparisons test. Adjusted p values are shown in graph. (F) Summary of effects of fin ablation on sensory neuron development. Micrographs in panels A-E were mirrored in either control or ablated samples for easier comparisons. All images are from rPec segments at st31. Scale bars, 50 µm. Original tiled images are shown in Figure S12. See also Figure S12.

We next examined whether markers downstream of *ntrk* signaling are affected by loss of the fins. Expression of *etv4* and *ret*, two genes known to be regulated by limb-derived signals in mice^76,77^, was markedly depleted in fin-ablated skate (Figure 7B,C, S12C,D). By contrast, expression of *runx3* was not diminished, but instead expanded throughout the DRG (Figure 7B,C).

Quantification revealed that the fraction of RUNX3^+^ISL1/2^+^ in pectoral DRG increased from 53% to 82% on the ablated side (Figure 7D). This expansion of *runx3* expression appears to be transient as analyses at later stages of development (st32) revealed restricted expression of *runx3* to DRG subtypes (data not shown). At these later stages, however, the number of DRG neurons is also markedly reduced, and the remaining neurons likely consist of DRG targeting axial tissues that are unaffected by fin ablation.

In addition to specifying DRG identities, target-derived cues play critical roles in shaping the central projections of primary sensory axons. We therefore examined how fin removal affects sensory projections within the SC, prior to neuronal loss. In fin-ablated embryos at st31, VGLUT1⁺ sensory terminals were largely confined to the dorsal-most layers of the spinal cord, with few projections extending ventral to LHX2⁺ interneurons (Figure 7E, S12E, F). To quantify these changes, we subdivided the dorsal spinal cord into three laminar bins and measured average VGLUT1⁺ integrated densities within each domain (Figure 7E). In fin-ablated embryos, VGLUT1 signal was significantly reduced across all layers, with the greatest reduction observed in deeper spinal cord regions.

Together, these results demonstrate that target-derived cues regulate both the molecular specification and fidelity of dorsal sensory afferent targeting in *Leucoraja* (Figure 7F). Because sensory neurons ultimately degenerate after fin ablation, we cannot exclude the possibility that peripheral injury or compromised sensory neuron integrity contributes to the observed changes in gene expression and central projections. Nevertheless, these findings are consistent with a model in which target-dependent coordination of neuronal specification and connectivity is an ancestral feature of vertebrate somatosensory systems.

## Discussion

The vertebrate somatosensory system transforms environmental signals into patterns of neural activity that encode internal representations of the external world. In mammals, somatosensory neuronal diversity and connectivity are strongly influenced by target-derived cues, a mechanistic divergence from the spatiotemporal patterning systems that predominate in most animal nervous systems^3,15,17,78^. Analysis of the little skate reveals that reliance on retrograde signaling is deeply conserved, even as the molecular programs implementing neuronal identity have diverged.

Conserved circuit architecture is thus compatible with malleability in receptor usage and fate-determinant deployment, providing a framework for understanding how vertebrate somatosensory systems maintain stable organizational logic while allowing evolutionary innovation in specification programs.

### Origins of diversity and topographic organization in somatosensory circuits

Our findings demonstrate that spinal and DRG neurons in *Leucoraja* exhibit molecular diversity. Profiling of spinal neurons indicates conserved cardinal classes of dorsal neurons, extending previous observations in skate ventral locomotor spinal circuits^40^. Spatial transcriptomic analyses further uncovered a laminar organization within the dorsal spinal cord, with distinct layers defined by the expression of fate determinants. Specifically, the most superficial dorsal layers are characterized by expression of *lmx1a*, *lmx1b*, and *tlx3b*—markers associated with nociceptive circuits in mammals, whereas deeper dorsal populations express *lhx2*, *lhx9*, and *barhl2*. These major classes can be further subdivided by TF expression, in line with recent molecular and spatial transcriptomic studies revealing extensive molecular diversity in the murine dorsal spinal _cord51,52._

We identified DRG neurons expressing core molecular features indicative of mechanosensory and nociceptive sensory classes. At a broad level, DRG neurons in skate can be classified as either mechanosensory-like (*ntrk2a*^+^, *piezo2*^+^) or nociceptor-like (*ntrk1/3*^+^, *trp*^+^, *tac1*^+^). Within these classes, further diversification is achieved through combinatorial expression of transcription factors and functional determinants. Notably, skate DRG neurons express at least one member of the four main classes of *trp* channels^79^ (*trpv1*, *trpa1*, *trpm2*, *trpc4*). Moreover, multiple transcription factors are expressed in both DRG neurons and their spinal targets, raising the possibility that shared transcriptional programs contribute to wiring specificity between connected neuronal populations.

Despite several conserved features, we uncovered striking evolutionary divergence in the deployment of SN fate determinants. In skate, *runx3* is co-expressed with *ntrk1* and *ntrk3,* a combination that is incompatible with nociceptor specification in mammals^80,81^. In mice, ectopic expression of *ntrk3* in *ntrk1^+^*neurons disrupts nociceptor identity and promotes acquisition of proprioceptor molecular features^81^. Our findings conceptually parallel studies of sensory neuron development in zebrafish^82^. In zebrafish *runx3* functions in trigeminal nociceptor specification^82^, whereas in mammals *runx3* marks trigeminal mechanoreceptors^83^. Moreover, several tetrapod SN fate determinants, including *etv4*, *meis2*, and multiple *hox* genes, show divergent class-specific deployment patterns between rodents and birds^61,63,76,84,85^. Collectively, these findings indicate that while laminar organization and class-based circuit architecture are conserved, the molecular programs specifying sensory neuron identity have undergone substantial evolutionary diversification. Whether the “mixed” molecular identities of skate sensory neurons represent retention of an ancestral program or are a lineage-specific innovation remains to be determined.

### Target-dependent and -independent mechanisms of somatosensory neuron development

In mammals, early-born DRG neurons initially express several fate determinants broadly that are progressively restricted to specific subtypes during circuit maturation^5,86,87^. Our fin ablation experiments suggest that a stepwise resolution of molecular identity represents an ancestral mechanism of sensory neuron specification. Following fin removal, and prior to neuronal loss, skate DRG neurons failed to restrict *runx3* expression to *ntrk1/3*⁺ populations and did not activate key developmental genes such as *etv4* and *ret*. Disruption of these molecular programs was accompanied by defects in central connectivity, with sensory axons largely confined to superficial layers and an overall reduction in spinal innervation. Although some innervation defects may reflect impaired sensory neuron integrity, the coordinated disruption of molecular identity and central projection patterns support an ancestral role for retrograde signaling in coupling sensory neuron differentiation with circuit assembly.

We further identified SMAD phosphorylation as an early indicator of DRG neurons responding to target-derived cues. In mammals, BMP signaling plays distinct roles in sensory neuron development along the rostrocaudal axis. In the trigeminal system, BMPs contribute to SN somatotopic organization^74^, whereas in DRG neurons SMAD proteins integrate BMP and neurotrophin signaling^75^. Although SMAD1 is dispensable for DRG survival and subtype specification in mice, it is required for proper terminal branching at peripheral targets^75^. These findings raise the possibility that integration of BMP and neurotrophin signaling through SMAD pathways represents an evolutionarily conserved mechanism linking target-derived cues to SN maturation and connectivity.

Our results highlight a target-independent phase of sensory neuron specification. The segregation of *ntrk2^+^* and *ntrk1/3^+^* DRG populations occurred normally after fin ablation, indicating that initial SN class identity is established independently of peripheral targets. This intrinsic mechanism likely reflects differences in the birth order of SN classes. In mammals, DRG neurons are generated over two neurogenic waves distinguished by reliance on the TFs *neurogenin1* (*ngn1*) and *neurogenin2* (*ngn2*) ^88^. Early born *ngn2* lineages give rise predominantly to *ntrk2^+^* and *ntrk3^+^* neurons (mechanosensory and proprioceptive), while later born *ngn1* lineages generate *ntrk1^+^* neurons (nociceptors and thermoreceptors). These results support an ancestral biphasic model of DRG neurogenesis in which broad sensory classes acquire core aspects of their molecular identity prior to target innervation, with subsequent refinement and circuit integration mediated by target-derived signals.

### Evolution of nociception, mechanosensation, and proprioception

The extent to which cartilaginous fish experience pain remains debated^33,35,36^. Although pain is an affective state, distinct from the ability to detect potentially harmful stimuli, the absence of unmyelinated C-fibers in elasmobranchs has been cited as evidence for limited capacity for pain perception. Our findings indicate that skates possess circuit and molecular features consistent with nociceptive processing. Skate DRG neurons express multiple *trp* channels with caconical mammalian nociceptive markers including *tac1* and *calca*. These putative nociceptors resemble mammalian thinly myelinated Aδ-fibers that mediate rapid nociceptive signaling associated with “first pain” ^39^. It remains to be determined whether skate nociceptor-like sensory are polymodal or whether they instead comprise distinct populations specialized for mechanical and chemical nociception.

Our interest in elasmobranch somatosensory circuits was motivated by prior work demonstrating a conserved program of limb/fin innervation. In mammals, limb coordination relies on mechanosensory feedback from muscle-targeting proprioceptive sensory neurons (pSNs) ^89-93^.

We were intrigued to find two key determinants of mammalian pSNs, *runx3* and *ntrk3*, were expressed in skate DRG neurons. Paradoxically, however, *runx3*^+^ neurons in skate express nociceptor markers but lack mechanosensory molecular indicators such as *piezo2*^94^. Skate DRG neurons also lack *etv1* expression, a target-regulated determinant essential for pSN development^95,96^. We hypothesize that muscle-based proprioception may represent a later evolutionary innovation, potentially emerging in concert with terrestrial locomotion.

Despite an apparent absence of muscle proprioceptors, skates and other fish species likely regulate motor output using alternative mechanosensory feedback systems^97^. In adult zebrafish, proprioceptive signals are conveyed by intraspinal *piezo2*^+^ neurons that directly regulate locomotor output^98^. Although we did not detect spinal *piezo2*^+^ neurons at embryonic stages, we identified *piezo2*^+^ neurons distinguished by *etv4* and *onecut2* expression, suggesting parallel mechanosensory channels that separately contribute to touch perception and motor control. Given that cutaneous afferents can modulate motor output in mammals via deep dorsal spinal circuits^99-101^, we speculate these ancestral somatosensory pathways provided a foundation upon which dedicated proprioceptive circuits subsequently emerged.

Taken together, these findings indicate that conserved spinal circuit architectures and target-dependent maturation programs preserve somatosensory organization across vertebrates, even as the molecular strategies implementing sensory feedback—including receptor usage and fate determinant deployment—remain evolutionarily flexible. This decoupling of circuit architecture from molecular specification provides a general mechanism by which vertebrate sensorimotor systems accommodate evolutionary innovation while preserving functional organization.

### Limitations of the study

Our cell-type assignments and inferred homologies rely on transcriptional markers and spatial organization rather than direct functional readouts. Although marker expression and laminar positioning support conserved relationships between DRG subtypes and dorsal spinal neuronal classes, definitive linkage of molecularly-defined sensory types to modality-specific physiology and behavior will require functional assays. Because our studies focus on developmental pathways, our single cell analyses of embryonic DRG neurons likely underrepresent the full molecular diversity of mature sensory types. In addition, whether the spatial organization of the dorsal spinal cord is maintained in the adult remains to be determined. While our fin ablation demonstrates a requirement for peripheral cues in shaping DRG identities and central projection patterns, this manipulation likely alters multiple target-derived signals and tissue environments, complicating attribution to any single pathway. Future work combining targeted perturbations with electrophysiology, activity imaging, and behavioral analyses will be essential to connect molecular identities to circuit function.

## Resource availability

### Lead contact

Requests for further information and resources should be directed to the lead contact, Jeremy S. Dasen

## Materials availability

All new materials reported in this manuscript are available from the lead contact upon request.

## Data and code availability

Raw and processed for snRNAseq (accession number GSE338711), bulk RNAseq (accession number GSE334356), and 10X Xenium spatial transcriptomics (accession number GSE341169) are available through GEO.

## Acknowledgments

This manuscript is dedicated to the memory of Heekyung Jung, whose work laid the foundation for the concepts explored in this study. We thank Sarah Pfennig, Theresa Steele, Annie Vonasek, and Natalie Williams for technical assistance. This work was supported by NINDS R35NS116858 to JSD, NINDS T32 NS086750 to PWF, Belfer Neurodegeneration Consortium from MD Anderson, the Carol and Gene Ludwig Family Foundation, the National Multiple Sclerosis Society, and the National Eye Institute (R01EY033353) to SAL.

## Author contributions

HL and JSD designed the experiments, prepared figures, and wrote the manuscript. HL, PWF and SAL assisted in designing, performing, and analyzing PIPseq experiments. E S-F and AEC performed and analyzed fin ablation and histological experiments. HDS, HK, and MB analyzed and prepared figures for bulk RNAseq experiments. KR, SS, MA, and CL assisted in designing, performing, and analyzing Xenium spatial transcriptomics.

## Declaration of generative AI and AI-assisted technologies in the manuscript preparation process

During the preparation of this work JSD and HL used ChatGTP to correct grammar, identify typos, and assist in RNAseq analyses. After using this tool, the authors reviewed and edited the content and take full responsibility for the content of the publication.

## Supplemental information

Document S1. Supplemental Figures S1–S12

Table S1. Selection of genes for 300 gene custom Xenium panel, related to Figures 1-5.

Table S2. Summary of results from 300 gene panel in Xenium data, related to Figures 1-5.

Table S3. Analysis of transcription factor expression in Xenium data, related to Figure 5.

Table S4. Analyses of SC and DRG snRNAseq data

Worksheet 1. SC cluster statistics.

Worksheet 2. SC top ten genes in each cluster by percent difference.

Worksheet 3. SC top ten genes in each cluster by fold change.

Worksheet 4. DRG neuron (subDRG) cluster statistics.

Worksheet 5. DRG neuron (subDRG) top ten genes by percent difference.

Worksheet 6. DRG neuron (subDRG) top ten genes by fold change.

## STAR Methods

### Biological Samples

#### Little skate (Leucoraja erinacea)

Little skate embryos at desired developmental stages were obtained from the Marine Resources Department at the Marine Biological Laboratory (Woods Hole, MA). Egg cases were maintained in artificial seawater (Instant Ocean, Aquarium Systems) at 16°C. Embryos were anesthetized in seawater containing MS-222 (tricaine methanesulfonate; Sigma-Aldrich) prior to fin ablations and tissue collections. For fin ablations embryos were placed in a Sylgaurd plate containing MS-222 with a small hole cut in the center to hold the egg yolk. The fin bud was removed using sharp scissors and returned to Instant Ocean and incubated 14-28 days at room temperature.

#### Chick

Fertilized chicken eggs (AVS-BIO, formerly Charles River Laboratories) were incubated at 39.5°C in a humidified incubator until the desired stage. Unilateral limb bud ablation was performed at Hamburger–Hamilton (HH) stages 16–18. Limb buds were removed by removing the top of the eggshell, injecting Pelikan ink under the embryo, and removing the limb using sharp forceps. Following ablation, the egg was sealed with Parafilm and incubated for 3-4 days prior to fixation.

All animal protocols used were approved by the Institutional Animal Care and Use Committee (IACUC) at NYU Langone Health.

### Fixation and sectioning

Anesthetized embryos were eviscerated and fixed in 4% paraformaldehyde (PFA) at 4°C for 1-2 hours, then washed 3-4 times in PBS (15-30 minutes per wash). For cryoprotection, embryos were incubated overnight in 30% sucrose at 4°C, embedded in Tissue-Tek OCT compound (Sakura Finetek), and then frozen at -80°C. They were subsequently cryosectioned at a thickness of 16 μm for antibody staining and HCR.

### Immunohistochemistry

Sections were washed in PBS to remove OCT compound, then blocked with 1% bovine serum albumin (BSA) in PBST (PBS with 0.1% Triton X-100) in humidified trays. Primary antibodies diluted in PBST with 0.1% BSA were applied and incubated overnight at 4°C. Sections were washed three times in PBST (5 minutes each), then incubated with secondary antibodies (1:1000 in PBST, filtered through a 0.2 μm syringe filter) for 1 hour at room temperature. Following three washes in PBST and one in PBS, coverslips were mounted using Vectashield (Vector Laboratories). Confocal images were acquired on a Zeiss LSM700 microscope.

The antibodies used are as follows: Primary antibodies: Guinea pig anti-RUNX3 (1:16K, Dasen lab stock), guinea pig anti-VGLUT1 (1:32K), mouse anti-ISL1/2 (1:50, Developmental Studies), mouse anti-LHX2 (1:100, Developmental Studies), rabbit anti-pSMAD (1:5000, Jessell lab), rabbit anti-FOXP1. Secondary antibodies were used at 1:1000 and are listed in the Key resource table.

Antibodies against *Leucoraja* proteins were generated in guinea pigs using the following peptide sequences at Labcorp Early Development Laboratories Inc. (formerly Covance) - VGLUT1-N: MEIRKEQLRKLAGDGLGRC, VGLUT1-C: CEEGKDVYMYGSAEDRDLP, RUNX3: EPVSLTQPNPPGLSRIRC. Antibody specificity was examined through direct comparison of antibody staining with mRNA expression patterns determined by HCR.

### DIG in situ hybridization

Tissue sections were air-dried for 10-15 minutes at room temperature and fixed in 4% PFA for 10 minutes. After two PBS washes (3 minutes each), sections were treated with Proteinase K (1 μg/mL) for 5 minutes, re-fixed in 4% PFA, washed, and incubated in triethanolamine for 10 minutes to block positive charges. Sections were washed three times in PBS, then blocked for 2–3 hours in hybridization solution (50% formamide, 5× SSC, 5× Denhardt’s solution, 0.2 mg/mL yeast RNA, 0.1 mg/mL salmon sperm DNA). Blocking solution was replaced with hybridization solution containing 100 ng of DIG-labeled antisense probe (100 μL per slide), and sections were incubated overnight at 72°C.

Following hybridization, slides were transferred to 5× SSC at 72°C for 20 minutes to remove coverslips, then washed in 0.2× SSC for 1 hour at 72°C. Sections were equilibrated in buffer B1 (0.1 M Tris pH 7.5, 150 mM NaCl) for 5 minutes at room temperature, then blocked with 10% heat-inactivated goat serum in B1. Blocking solution was replaced with anti-DIG-AP antibody (1:5000; Roche) in B1 containing 1% heat-inactivated goat serum, and sections were incubated overnight at 4°C in a humidified chamber. Sections were washed twice in B1 (5 minutes each), equilibrated in buffer B3 (0.1 M Tris pH 9.5, 100 mM NaCl, 50 mM MgCl2) for 5 minutes, and developed in B3 containing BCIP and NBT (3.5 μL/mL each) for 12–48 hours. After color development, sections were washed in ddH₂O and mounted in Glycergel (Agilent).

### Hybridization chain reaction (HCR) RNA-FISH

HCR was performed on fresh-frozen tissue sections following the third-generation protocol (v3.0) from Molecular Instruments^102^. DNA probe sets, HCR amplifiers, and associated buffers were obtained from Molecular Technologies (Beckman Institute, Caltech). Custom probes were synthesized by Integrated DNA Technologies using oPools Oligo Pools. Slides were mounted using 110 µl Vectashield (Vector Laboratories), and confocal images were acquired on a Zeiss LSM700 microscope. Probes with indicated amplifier sequences were as follows: B1: *runx3*; B2: *tbx3b, pou4f3, etv4, skor2, tlx3, hoxd1, onecut2, pou4f1, shox2, six1, piezo2, slc17a6, slc17a7, ntrk3, ret, ntrk1, ntrk2a.* Probe sequences are available from the Lead author upon request.

### Bulk RNA-seq library preparation

Spinal cord and dorsal root ganglia (DRG) were dissected from st32 *Leucoraja erinacea* embryos and stored separately in TRIzol reagent (Invitrogen) at −80°C. Samples were homogenized by trituration through 18-gauge (20×) and 25-gauge (10×) needles. Following addition of 100 μL chloroform, samples were incubated on ice for 5 minutes and centrifuged at 12,000 rpm for 15 minutes at 4°C. The aqueous layer was transferred to a new tube, mixed with 250 μL isopropanol, incubated on ice for 10 minutes, and centrifuged at 12,000 rpm for 10 minutes at 4°C. RNA pellets were washed twice with 500 μL 75% ethanol (7,000 rpm, 5 minutes, 4°C), air-dried, and resuspended in 25 μL EB buffer. Samples were further purified using the RNA MinElute Cleanup Kit (Qiagen) to remove DNA and small RNA contamination. Stranded RNA-seq libraries were prepared with polyA selection by the NYU Langone Genome Technology Center (RRID: SCR_017929) and sequenced on an Illumina NovaSeq 6000 using an SP flow cell (300 cycles, v1.5 chemistry).

Raw paired-end sequencing data underwent quality control and adapter removal using Trimmomatic^103^. Adapter trimming was performed based on the specific barcode sequences for each sample: DRG815 (TAAGATTA), DRG816 (ACGTCCTG), DRG818 (GTCAGTAC), SC815 (AGGATAGG), SC816 (TCAGAGCC), and SC818 (CTTCGCCT). The cleaned paired-end reads were then aligned to the *Leucoraja erinacea* reference genome (Assembly: ASM2654585v1) using HISAT2 (version 2.2.0) ^104^. Following alignment, the resulting BAM files were sorted and indexed using Samtools^105^. Gene expression levels were quantified by counting fragments mapping to exons using featureCounts^106^, generating a raw count matrix for downstream analysis.

### Bulk RNAseq differential expression analysis

To identify differentially expressed genes (DEGs) between SC and DRG tissues, we performed differential expression analysis using the edgeR package (version 4.4) in R^107^. The raw read counts were normalized using the Trimmed Mean of M-values (TMM) method to account for library size differences. Statistical significance was tested based on a negative binomial distribution model.

The results of the differential expression analysis were visualized using a volcano plot (Figure S2A). Significant DEGs were defined based on stringent criteria: an adjusted P-value (FDR) < 0.01 and an absolute log2 fold change (|logFC|) > 3. Genes upregulated in the SC are shown in red, while genes upregulated in the DRG are shown in blue. To further characterize tissue-specific signatures, expression profiles of transcription factors and selected DRG-enriched genes were analyzed (Figure S2B and S2C). For these visualizations, gene expression levels were quantified as Reads Per Kilobase per Million mapped reads (RPKM). The data were visualized using dot plots, where the dot size represents the relative count, and the color intensity corresponds to the log-transformed RPKM values. All visualizations were generated using the ggplot2 package^108^ in R.

### Nuclear isolation

Spinal cords and dorsal root ganglia (DRG) were dissected from st31 and st32 *Leucoraja erinacea* embryos (Marine Biological Laboratory, Woods Hole, MA) in seawater and collected in 500 μL filtered sEBSS (∼45 mL EBSS supplemented with 1 g urea, 0.7 g NaCl, and 0.2 g TMAO). Samples were centrifuged at 300 × g for 4 minutes at 4°C, supernatant was removed, and pellets were snap-frozen on dry ice and stored at −80°C.

Nuclei were isolated from spinal cords (6 embryos) and individually dissected DRG (14 embryos). Frozen samples were mechanically homogenized in 2 mL cold EZ Lysis Buffer (Sigma) using a Dounce homogenizer (20 strokes loose pestle, 20 strokes tight pestle). The homogenizer was rinsed with 1 mL EZ Lysis Buffer, samples were brought to 5 mL total volume and incubated on ice for 5 minutes. Samples were centrifuged at 500 × g for 5 minutes at 4°C, resuspended in EZ Lysis Buffer, and the lysis step was repeated. Pellets were resuspended in nuclei suspension buffer (PBS containing 0.1 mg/mL BSA and 0.2 U/μL RNase inhibitor), centrifuged, resuspended in ∼1 mL nuclei suspension buffer, filtered through a 40 μm cell strainer, centrifuged again, and resuspended in 20 μL cell suspension buffer.

### Single-cell and single-nucleus capture via PIPs

Approximately 45,000 nuclei (DRG) and 60,000 nuclei (SC) were loaded for library preparation following the manufacturer’s instructions. Single cells and nuclei were partitioned into pre-templated instant partitions (PIPs) following the manufacturer’s instructions (Fluent BioSciences, now Illumina). Briefly, 40,000–60,000 nuclei were combined with template beads (thawed on ice), mixed gently by pipetting, and then vortexed for 175 seconds to isolate single cells. Reverse transcription and library prep were then performed according to the manufacturer’s protocol. Samples were sequenced on an Illumina NovaSeq X (target: 25,000 reads/nucleus; achieved: 66,581 reads/nucleus for DRG, 4,687 reads/nucleus for SC).

### Skate bioinformatics

The chromosome-level genome assembly and annotated transcriptome of the little skate^109^ were accessed from NCBI (RefSeq: GCF_028641065.1). Chromosome names in the genome were edited to match the transcriptome chromosome names, then the edited .fna and .gtf files were used to make an indexed genome using STAR^110^. This indexed genome was used to process raw .fastq files via PIPSeeker (v2.1.4), a wrapper function for STARSolo. Droplet detection was performed using Cellbender. Quality control metrics for each nucleus included the number of detected genes (nFeature_RNA) and total UMI counts (nCount_RNA). Nuclei were excluded from further analysis based on the following cutoffs: n_gene_by_counts <2500, total_ counts <5000. Standard preprocessing and normalization were performed in Seurat (v.5.1.0), and data were integrated across samples. Total number of nuclei analyzed: DRG+SC: 22,274 (Post QC), 75,213 (Pre QC), SC: 15,697 (Post QC), 27,600 (Pre QC), DRG neurons: 6,577 (Post QC), 47,553 (Pre QC)

Marker genes for each cluster and related differential expression (DE) analyses were calculated using the Wilcoxon rank-sum algorithm implemented in Seurat. All statistical tests used are highlighted in the legend of each figure. No data were excluded from the analyses, except when performing quality control filtering as discussed above. For analyses of DRG, nuclei were first clustered in Seurat without reference to marker genes (FindNeighbors, dims = 1:30; FindClusters, resolution = 0.5), yielding the ten clusters in Figure S7A. Non-neuronal clusters were identified by cluster-restricted markers and excluded, and the three neuronal clusters (1, 2, and 3), expressing sensory-neuron markers (e.g., runx3, ntrk2a, ntrk3), were retained. These nuclei were then re-clustered in Seurat (FindNeighbors, dims = 1:30; FindClusters, resolution = 1.5), and non-sensory subclusters, identified by their separation in UMAP and marker expression, were removed; the retained subclusters were re-clustered once more (FindNeighbors, dims = 1:30; FindClusters, resolution = 1.5) to yield the seven neuronal clusters in Figure 3.

Marker gene expression informed only which clusters were carried forward at each step; it did not influence the selection of individual nuclei, clustering resolution, or cluster boundaries, all of which were determined by unsupervised clustering. Experiments were not randomized. The investigators were not blinded to allocation during experiments and outcome assessment. Gene names follow the *Leucoraja erinacea* genome annotation where available (e.g., *ntrk2a*). For clarity, genes annotated only with LOC identifiers are referred to by the names of their presumed mouse homologs. Corresponding LOC identifiers are provided in Tables S1 and S2.

### Xenium Spatial Transcriptomics

Spatial transcriptomic profiling was performed using the 10x Genomics Xenium In Situ platform at the Experimental Pathology Research Laboratory at NYU Langone Health. Skates were fixed in 4% PFA for 2.5 hours, washed in PBS 3-4 times, 30 minutes each, and incubated overnight in 30% sucrose. Samples were embedded in OCT, frozen on dry ice, and sectioned at 10 μm and placed directly onto room temperature Xenium slides. Sections were fixed, permeabilized, and hybridized with a Xenium Custom gene panel targeting 300 genes. Following probe hybridization, signal amplification and fluorescent detection were carried out on the Xenium Analyzer.

High-resolution imaging was used to detect individual RNA molecules in situ, and cell segmentation was performed using DAPI nuclear staining combined with membrane and cytoplasmic boundary staining using Xenium Multi-Tissue Stain Mix. Raw image processing, transcript decoding, and cell segmentation were conducted using the Xenium Onboard Analysis pipeline. Downstream quality control and analysis were performed using Xenium Explorer 4. Probes with low signal were excluded from further analysis. Spatial gene expression patterns were visualized and analyzed in the context of tissue morphology using Xenium Explorer 4.

### Analyses of vGlut1 in the dorsal spinal cord

For analyses of vGlut1 innervation density in the dorsal spinal cord, 16-bit tiled confocal images spanning control and fin ablated regions were collected using 10 µm optical sections with a 20X objective on a Zeiss LSM700 confocal microscope. Laser power settings were manually adjusted in each section to achieve sub-maximal signal intensities. Images were analyzed in FIJI, first by subtracting background using 50 µm rolling paraboloid. Three equal size regions of interest (ROIs) were applied to control and ablated spinal cord regions in each image. Intensity densities for each embryo were averaged from 6 sections per animal. Differences in intensities between ablated and control sections were calculated using 2-way ANOVA with Šídák’s multiple comparisons test.

