## Supplemental Figures for "Ancient Somatosensory Circuit Architectures Employ Flexible Molecular Strategies"

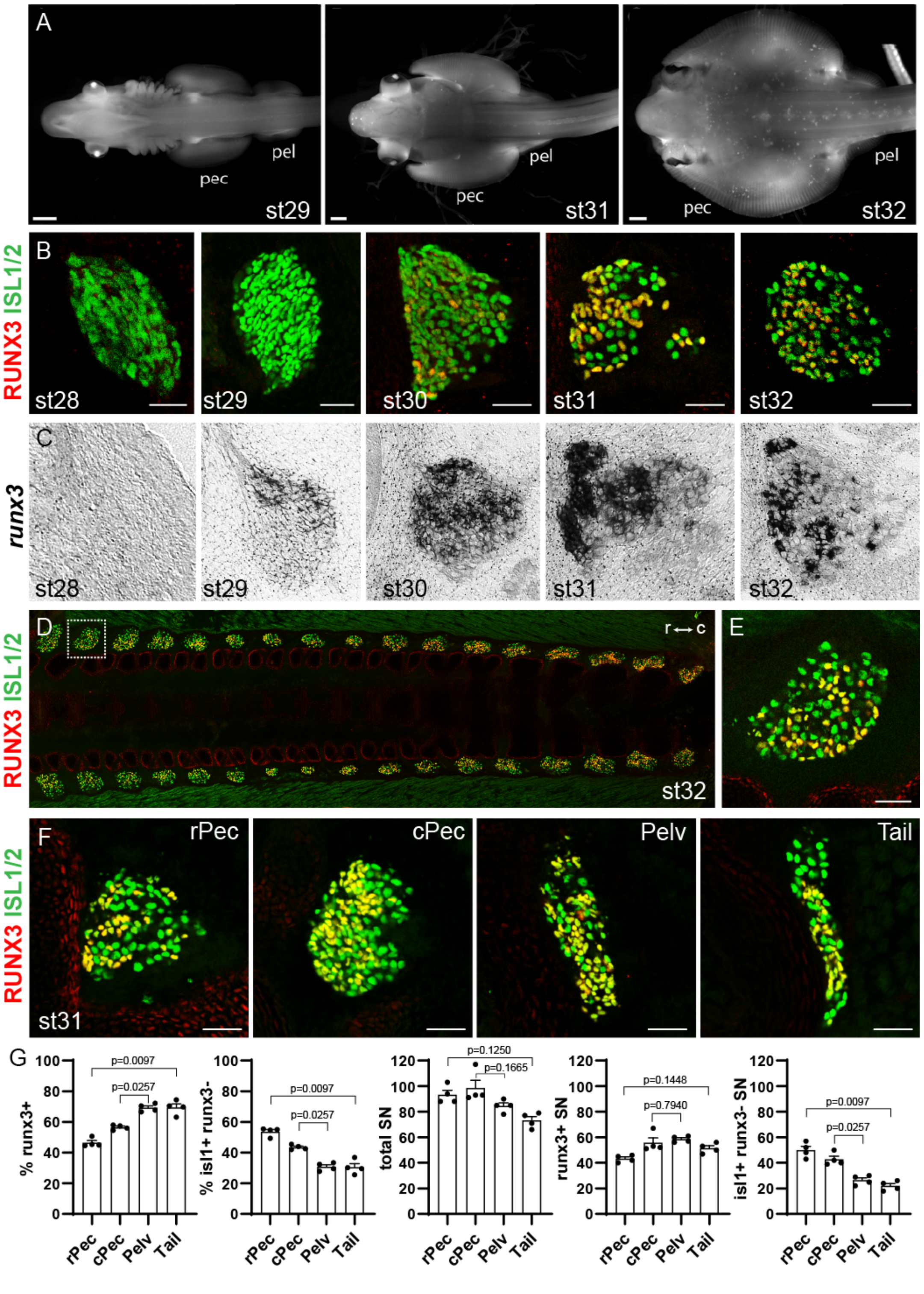


**Figure S1. Ontogeny of DRG sensory neuron development in *Leucoraja*.** (A) Development of midgestation little skate embryos at st29, st31, and st32. Modified from Gillis et al.[23] Scale bars, 1 mm. (B) Expression of RUNX3 and ISL1/2 proteins in DRG between st28 and st32 in rostral pectoral segments. Scale bars, 50 µm. (C) Expression of *runx3* mRNA in rPec segments between st28 and st32. DIG-labeled *runx3* anti-sense probe was used. (D) Top-down view of st32 skate DRG along rostral (r) - caudal (c) axis stained with RUNX3 and ISL1/2. (E) Expression of RUNX3 and ISL1 in region indicated in panel D. (F) Expression of RUNX3 and ISL1/2 at indicated segmental levels at st31. Scale bars, 50 µm. (G) Quantification of the percentages and number of DRG neurons expressing RUNX3 and ISL1 at indicated rostrocaudal levels, averaged from 4 embryos at st31 (3 DRG sections per embryo). Data are presented as mean ± SEM, with each point representing one embryo. Statistical significance for rPec vs. Tail and cPec vs. Pelv were determined using a one-way repeated-measures ANOVA with Geisser–Greenhouse correction, followed by Tukey's multiple-comparisons test. Exact adjusted *P* values are shown in the figure.

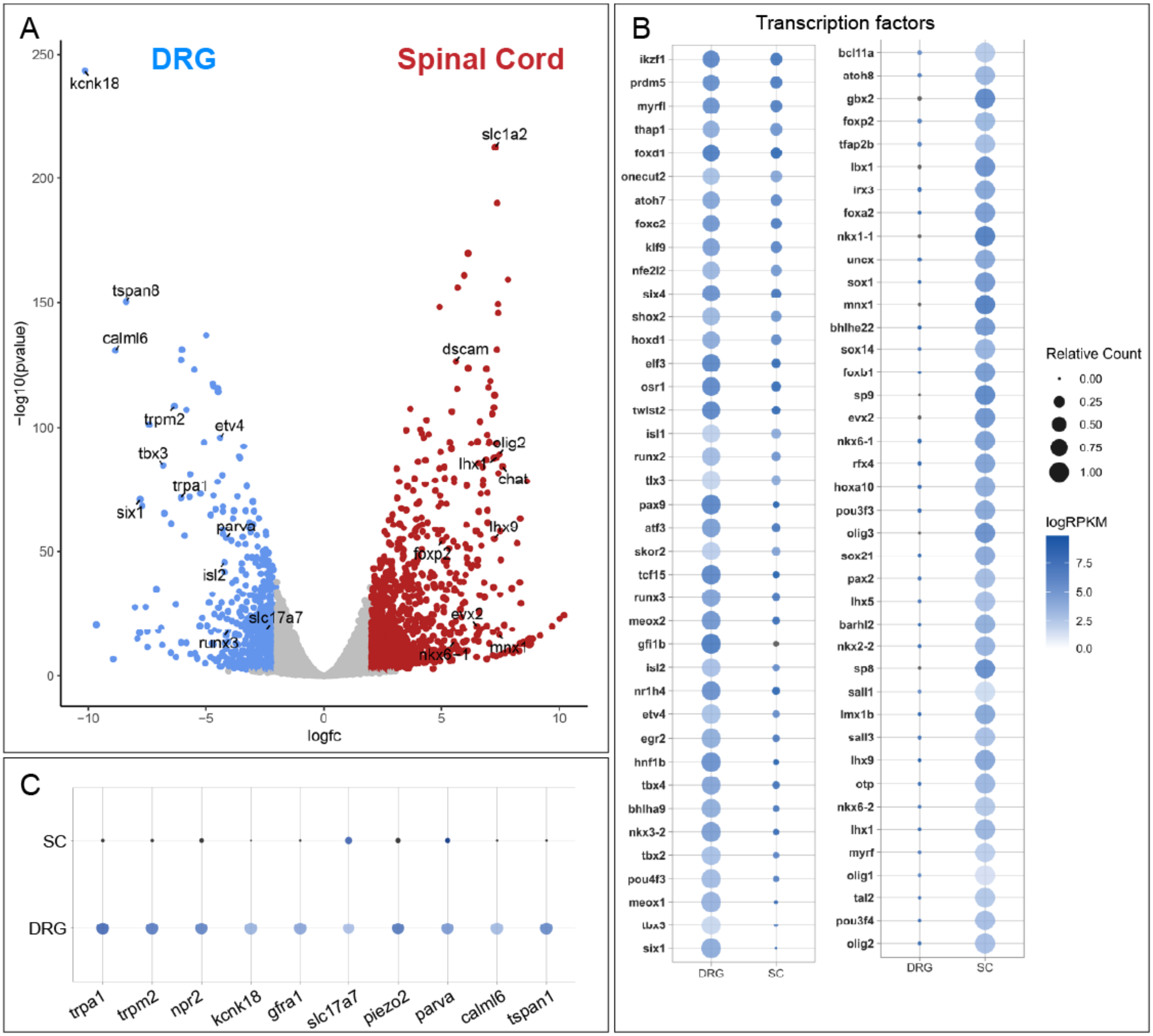


**Figure S2. Bulk RNAseq analysis of *Leucoraja* spinal cord and DRG tissue.** (A) Volcano plot of SC and DRG enriched genes. Genes that are conserved with mammals or showing highly differential expression between SC and DRG are highlighted. (B) Dot-plots of transcription factors enriched in either DRG (n=3) or SC (n=3) samples. (C) Dot plots of selected DRG-enriched genes.

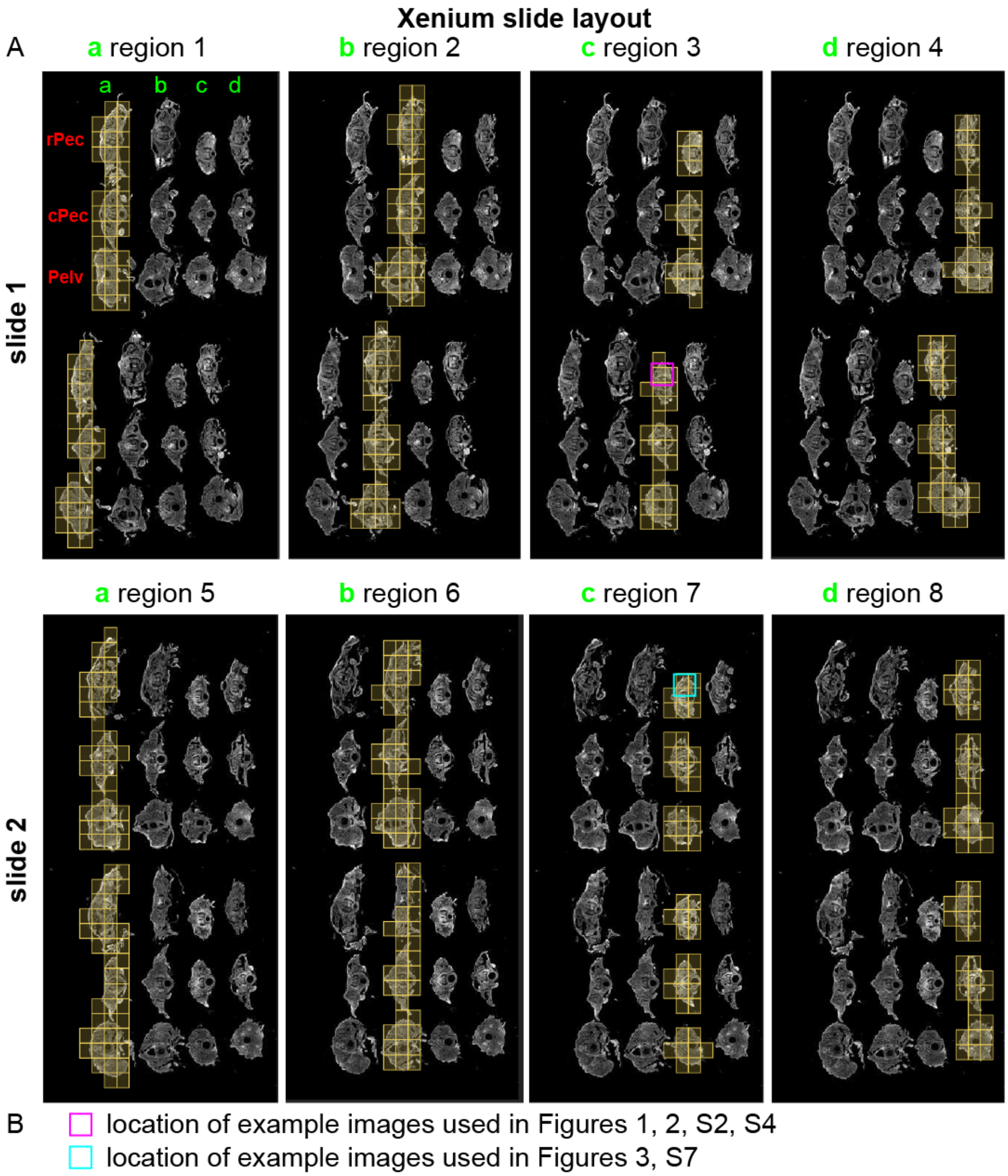


| embryo | stage at FBA | days incubated | FBA side | stage fixation | "embryo" Xen. File | notes |
|---|---|---|---|---|---|---|
| a | 28 | 28 | R | 32 | 1,5 | no fin regrowth |
| b | 28 | 28 | L | 32 | 2,6 | no regrowth - lost pelvic SC |
| c | 28+ | 20 | L | 31+ | 3,7 | no fin regrowth |
| d | 28+ | 20 | L | 31+ | 4,8 | small regrowth pec |

**Figure S3. Layout of samples analyzed using Xenium spatial transcriptomics.** (A) Overview image of sections mounted on two Xenium slides. Each slide contains two sections, each section having 4 embryos with rostral pectoral (rPec), caudal pectoral (cPec), and pelvic (Pelv) segments. Gold boxes indicate scanned regions. Pink and cyan boxes indicate the positions of example images used in the Figure. (B) Table indicating stage of embryos at collection, incubation times, and side of fin ablation (FBA). Fin ablation results were not used for molecular profiling DRG due to substantial neuronal loss at these late stages.

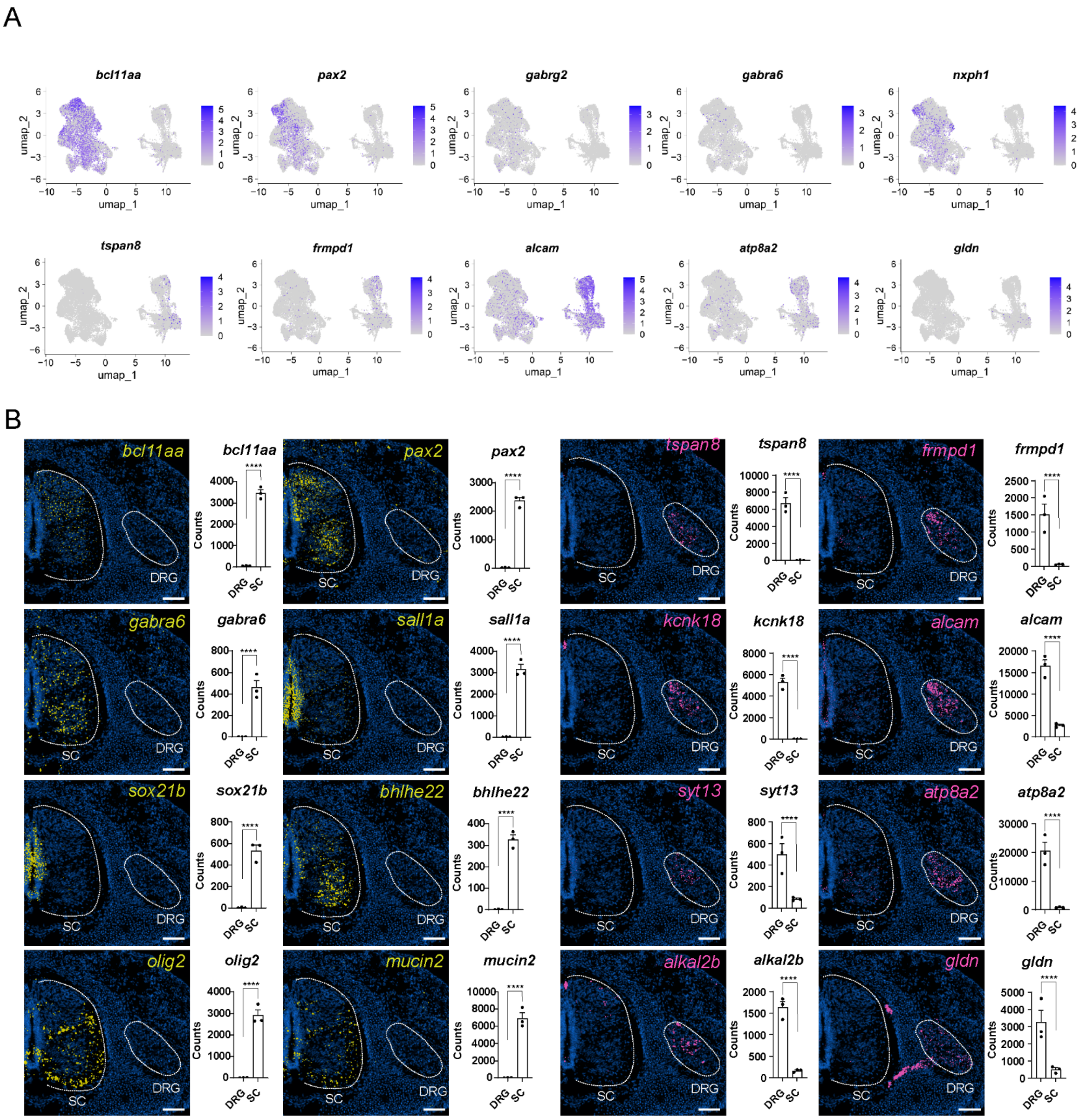


**Figure S4. Validation of candidate genes by spatial transcriptomics.** (A) Feature plots showing additional examples of SC- and DRG-enriched genes. Note detection of certain genes (e.g. *gabrg2*) is less robust by snRNAseq than using Xenium or bulk RNAseq. (B) Spatial transcriptomic analyses SC- and DRG-enriched genes. Images are shown from a single rostral pectoral section at st31+. Scale bars, 100 µm. Similar patterns were observed in multiple segments from n=4 embryos analyzed at st31+ (n=2) and st32 (n=2). Plots on right show mean ± SEM RPKM from three biological replicates. Statistical significance was determined from raw read counts using edgeR with TMM normalization and a negative-binomial model, with Benjamini–Hochberg correction for multiple testing. ****FDR < 0.0001.

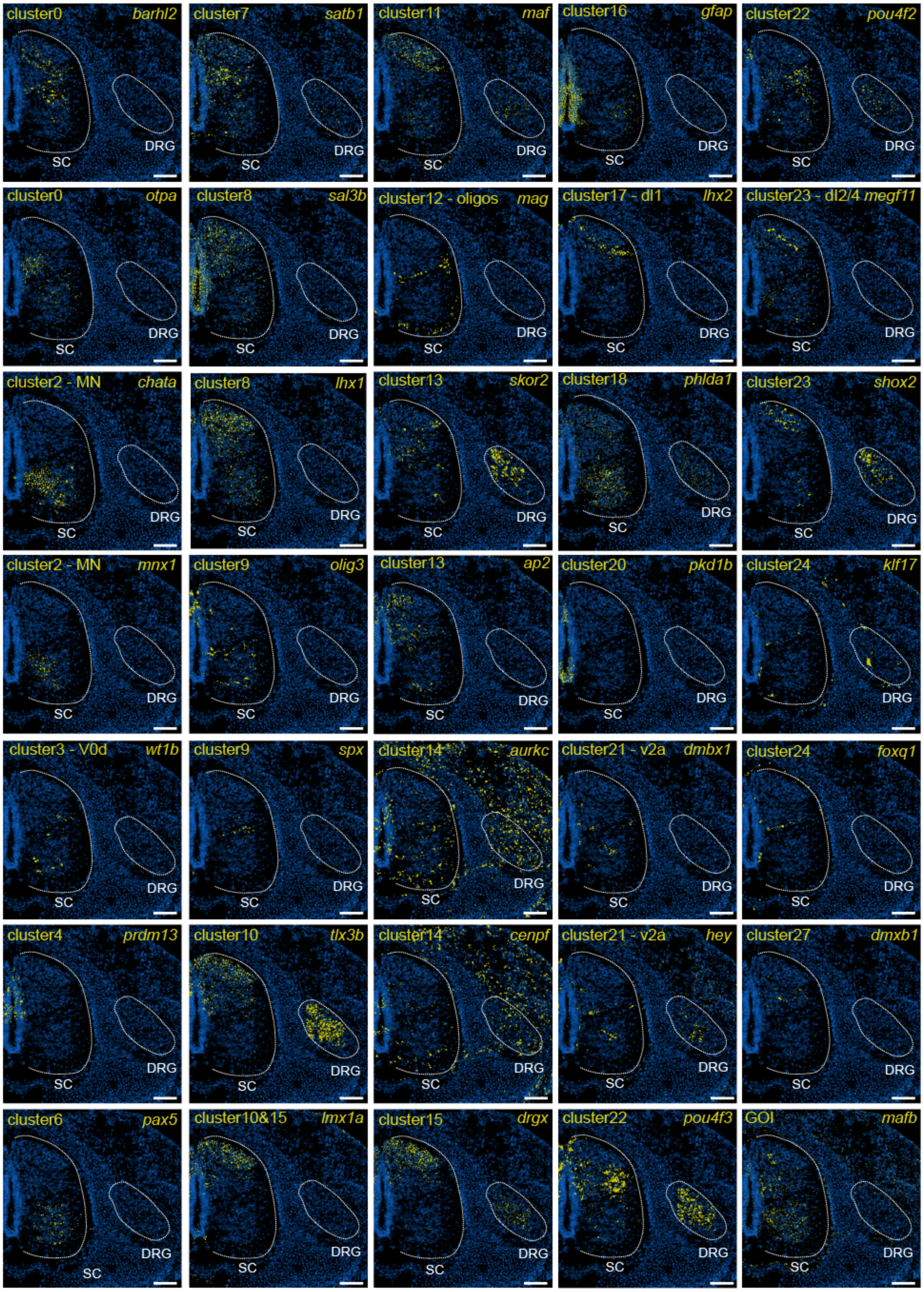


**Figure S5. Spatial transcriptomic analyses of enriched genes in spinal cord clusters.** Images of Xenium spatial transcriptomics of cluster-restricted genes in st31+ skate spinal cord. Images indicate cluster origin from UMAP plot shown in Figure 2A and a gene of interest (GOI), *mafb*. Scale bars, 100 µm. At least 2 genes were chosen from each cluster. Spinal cord (SC) and dorsal root ganglia (DRG) are outlined in each image. DAPI staining is shown in blue. Similar patterns were observed in multiple segments from n=4 embryos analyzed.

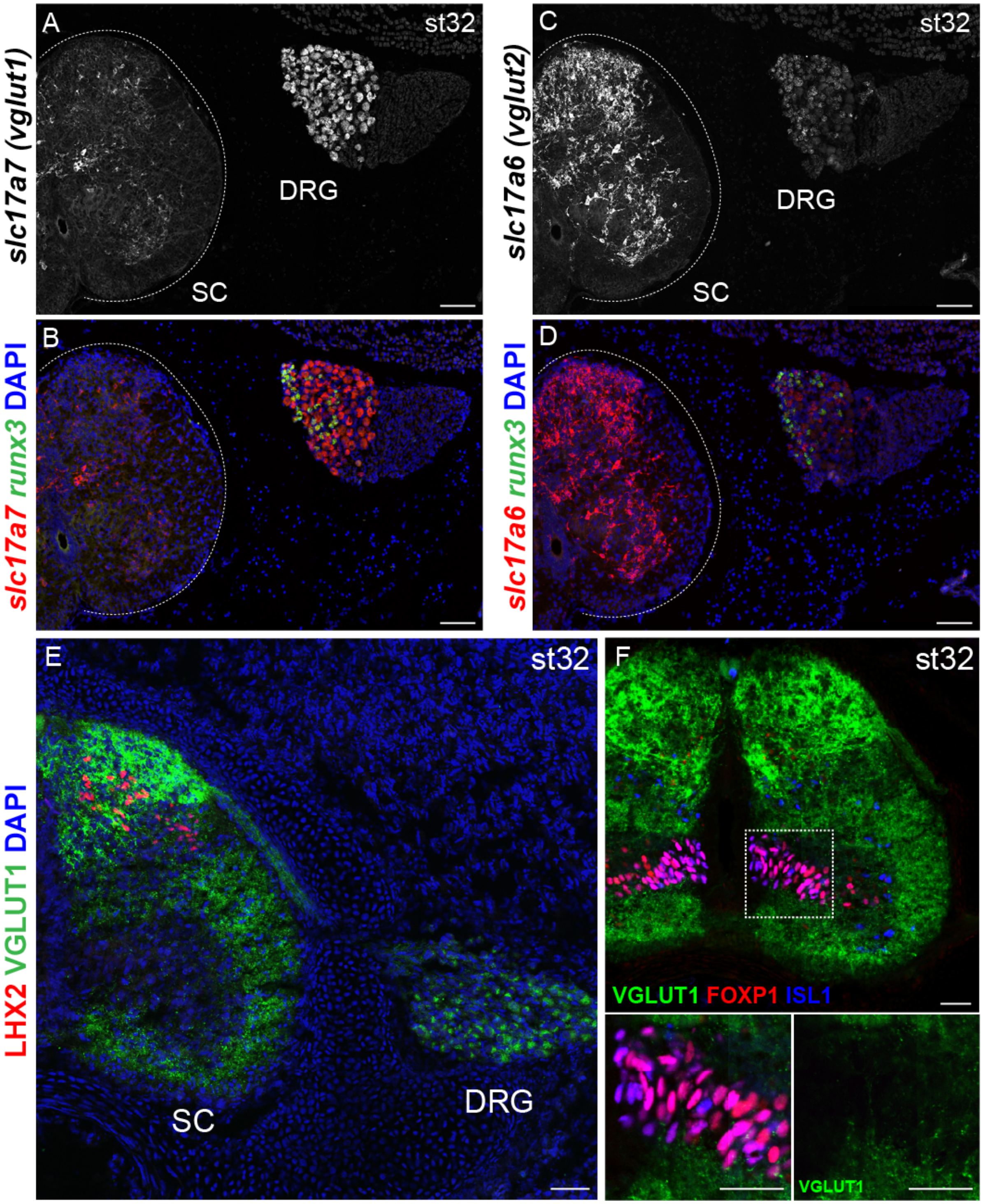


**Figure S6. Analyses of *vglut1* and *vglut2* expression in spinal cord and DRG of *Leucoraja*.** (A) HCR *in situ* hybridization showing expression of *slc17a7* (*vglut1*) in DRG and SC of rostral pectoral (rPec) segments at st32. Note *slc17a7* is detected in a small population of medial SC cells at this level. (B) Expression of *slc17a7* and *runx3*. (C) Expression of *slc17a6* (*vglut2*) in DRG and SC of rostral pectoral segments at st32. (D) Expression of *slc17a6* and *runx3*. In A-D there appears to be non-neuronal cells on the lateral side of the DRG. (E) Staining of VGLUT1 and LHX2 in SC and DRG. Note absence of VGLUT1 staining in the ventral spinal cord. DAPI is shown in blue. (F) Staining of VGLUT1, FOXP1, and ISL1 at rPec levels at st32. Inset shows undetectable VGLUT1 staining near FOXP1$^{+}$ fin motor neurons. Scale bars in A-D, 100 µm; E-F, 50 µm. Magnified images of panels B, D, and E are shown in the main Figure 2.

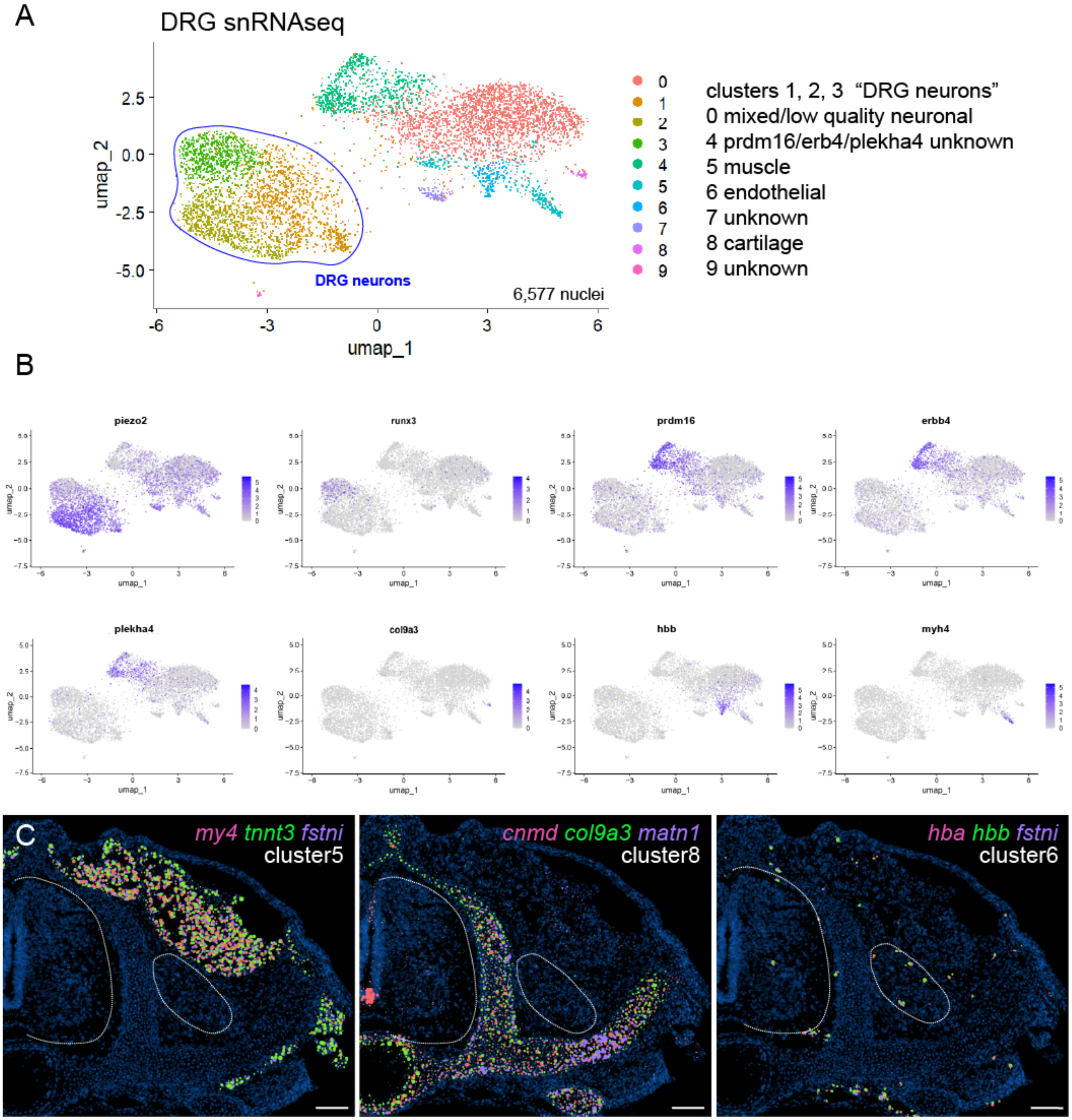


**Figure S7. Analyses of snRNAseq of DRG in *Leucoraja*.** (A) UMAP plot of DRG cells showing cells selected for DRG neuron analyses. Non-neural nuclei indicated on right. Clusters 1, 2, 3 were reclustered as "DRG neurons" based on expression of *runx3*, *ntrk2a,* and *ntrk3*, (B) Feature plots showing expression of selected genes. (C) Spatial transcriptomics of three genes from non-neural clusters showing expression in presumptive muscle (cluster5), cartilage (cluster8), and endothelial cells (cluster 6). Sections are from rPec levels of a st31+ embryo. DAPI is shown in blue. Scale bars, 100 µm.

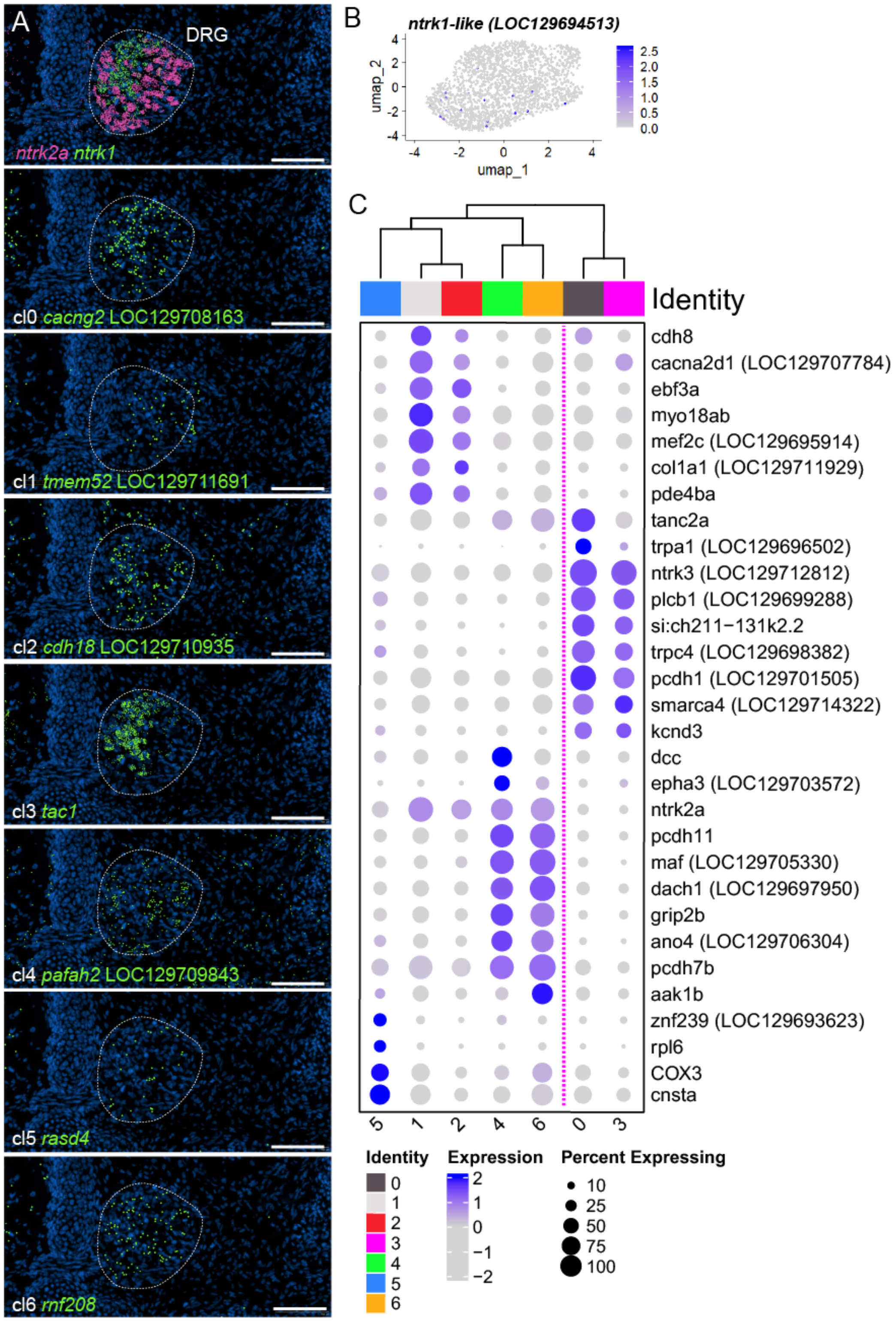


**Figure S8. Spatial transcriptomics of cluster-defining genes in DRG neurons.** (A) Example Xenium images showing expression of DRG neuron cluster-enriched genes from UMAP plot shown in Figure 3. Images are from a single rPec section and representative of n=4 st31+ and n=4 st32 embryos analyzed. *ntrk1* and *ntrk2a* expression are shown for reference. DAPI stain is shown in blue. Scale bars, 100 µm. (B) The *ntrk1*-like gene is not well detected in snRNAseq data. (C) Dot plot of selected genes in each of the seven DRG clusters. Cluster 5 may be non-neuronal. Magenta line shows separation of putative skate nociceptor and mechanosensory neurons.

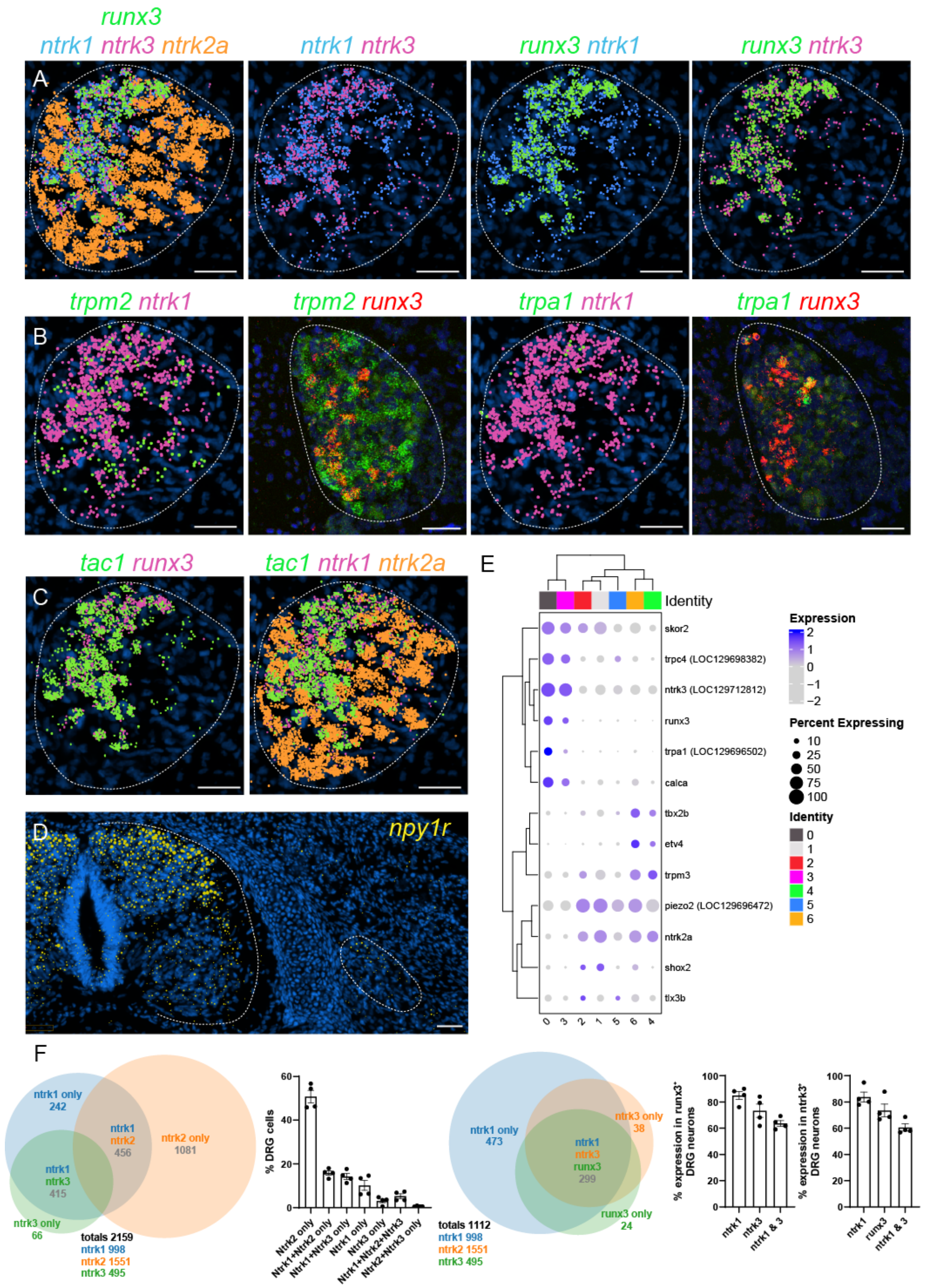


**Figure S9. *Ntrk* gene expression and sensory modality markers in *Leucoraja*.** (A) Expression of *runx3* in relation to *ntrk* genes. (B) Expression of *trpm2* and *trpa1* in DRG. *Trp* channel expression is shown by both Xenium (left panels) and HCR (right panels). (C) Expression of *tac1* in *runx3*$^{+}$ and *ntrk1*$^{+}$ neurons. (D) Expression of *npy1r* is enriched in the superficial dorsal horn. Images are representative of n=4 embryos analyzed at st31+ and st32. DAPI (blue) is shown in each panel. Scale bars, 50 µm. (E) Dot plots of selected DRG-enriched genes showing expression of ntrk, TFs, and modality-restricted genes in DRG neuron clusters. (F) Venn diagrams and percentages of neurons expressing *ntrk* genes (left panel) and *ntrk1*, *ntrk2a*, and *runx3* (right panels) from 4 embryos counted in Xenium data. Total cells counted shown in Venn diagrams. Percentages from each embryo (4 Pec sections/embryo) shown in graphs and are presented as mean ± SEM, with each point representing one embryo.

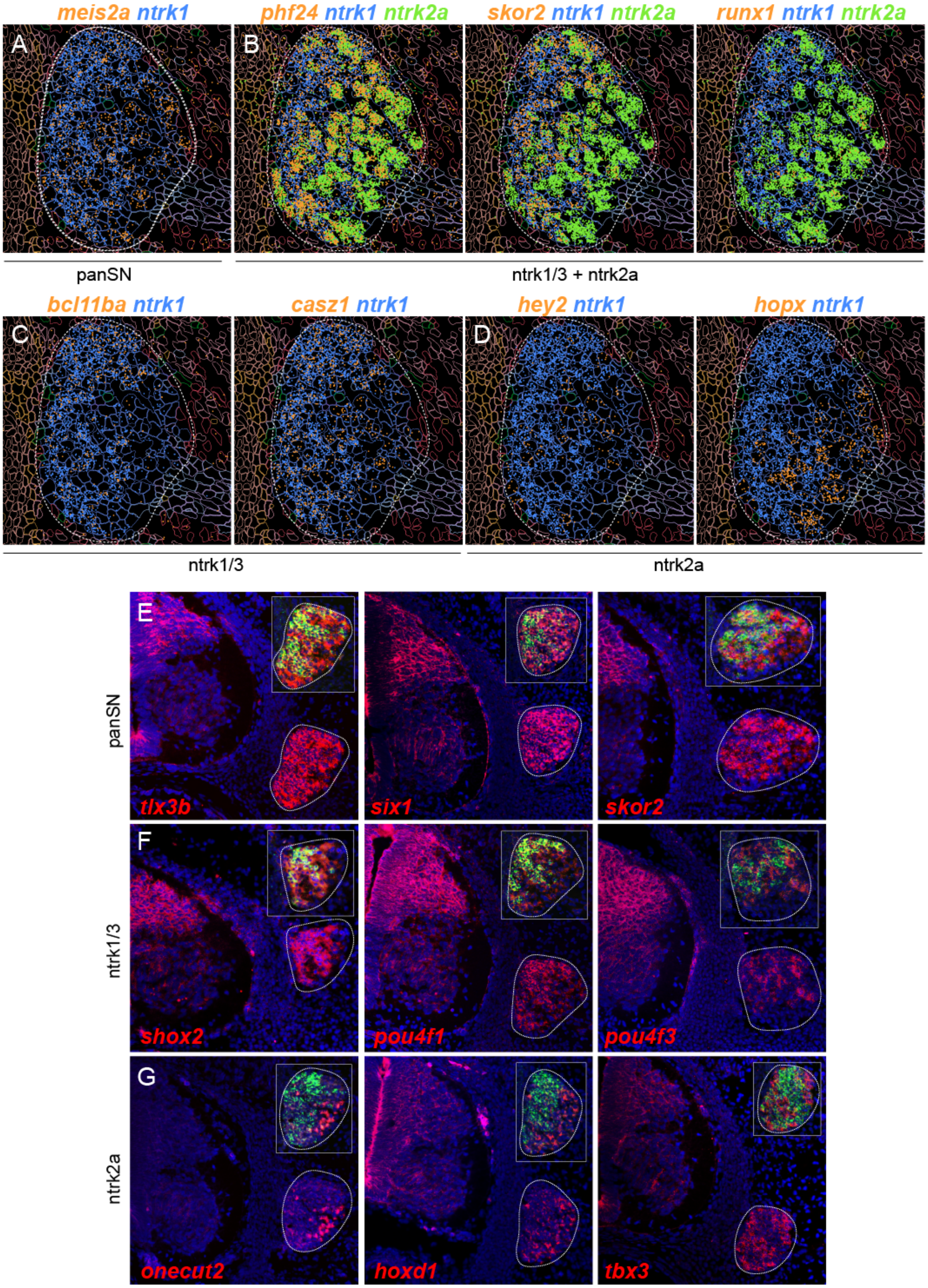


**Figure S10. Transcription factor expression in DRG neurons of *Leucoraja*.** (A) Expression of *meis2a* in DRG. (B) Expression of *phf24*, *skor2*, *runx1* are detected in subsets of both *ntrk1/3*+ and *ntrk2a*+ cells. (C) *bcl11ba* and *casz1* are detected in *ntrk1/3*+ cells. (D) *hey2* and *hopx2* are detected in *ntrk2a*+ cells. Scale bars, 50 µm. (E) Tiled confocal images showing further validation of TF expression by HCR *in situ* hybridization. Insets show expression of TF of interest (red) in relation to *runx3* (green).

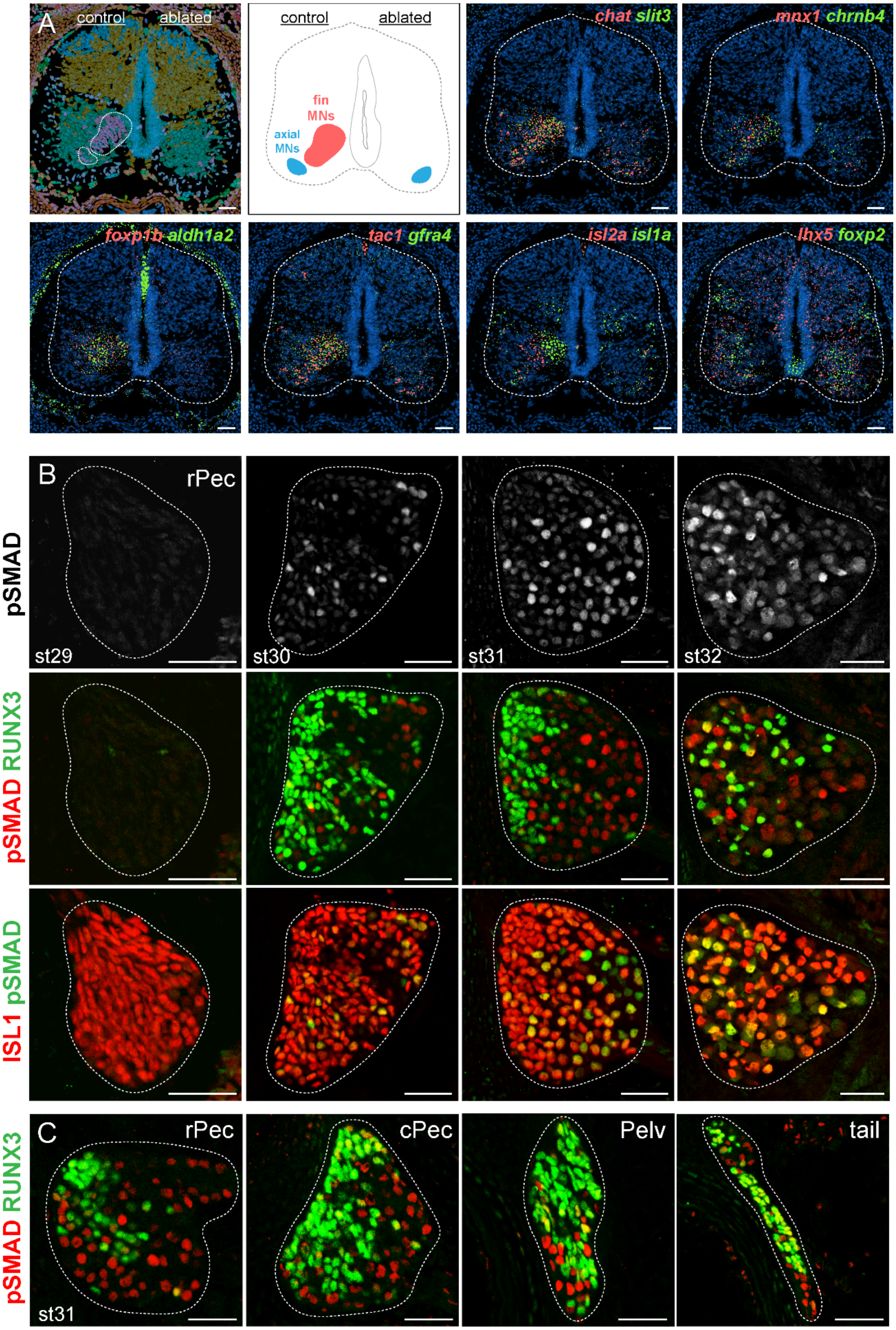


**Figure S11. Effect of fin bud ablation on motor and sensory development.** (A) Impact of fin bud ablation on motor neurons. Spatial transcriptomics of indicated genes showing reduced number and expression of MN-restricted genes on ablated side. Expression of *lhx5* and *foxp2*, two interneuron markers, is grossly unchanged. Images are from rPec levels, st32. (B) Expression of pSMAD in skate between st29 and st32 in rPec segments. SMAD phosphorylation is detected beginning at st30. (C) Expression of pSMAD at indicated segmental levels at st31. Scale bars, 50 μm.

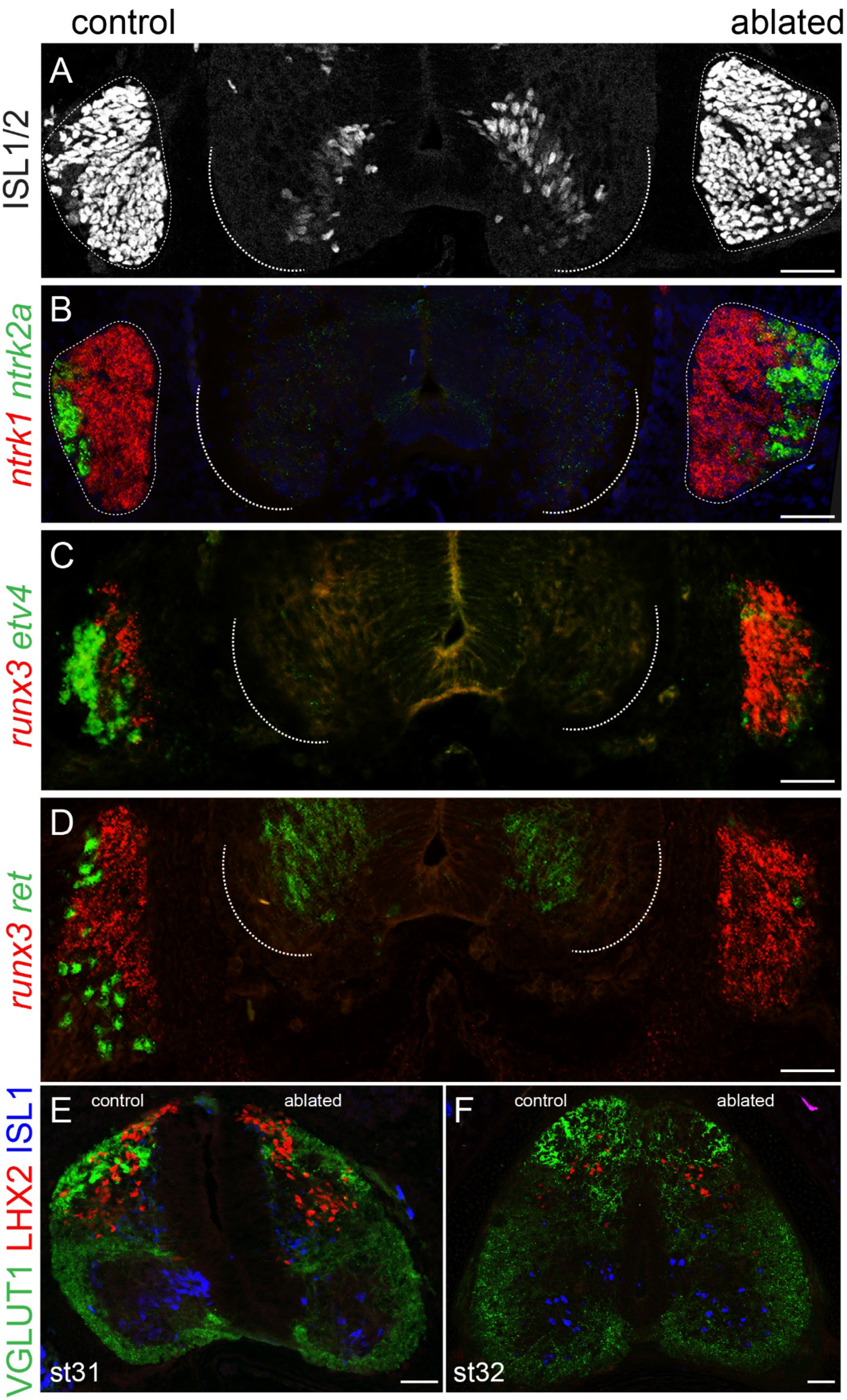


**Figure S12. Role of fin-derived cues in sensory development and connectivity.** (A-D) Tiled confocal images of a single section from control and fin-ablated embryos showing changes in DRG gene expression. (E-F) Tiled confocal images showing reduced VGLUT1 terminal density in fin-ablated embryos at st31 and st32 in rPec segments. Similar results were obtained from n=6 embryos analyzed at st31 and n=4 embryos at st32. Scale bars, 50 μm.
